# Cells as Single-Molecule Protein Proximity Networks

**DOI:** 10.1101/2025.06.19.660329

**Authors:** Filip Karlsson, Michele Simonetti, Christina Galonska, Max Karlsson, Hanna van Ooijen, Divya Thiagarajan, Tomasz Kallas, Maud Schweitzer, Ludvig Larsson, Vincent van Hoef, Pouria Tajvar, Johan Dahlberg, Florian De Temmerman, Louise Leijonancker, Vanessa Trombin, Sylvain Geny, Rikard Forlin, Erika Negrini, Beijing Wu, Liu Xi, Stefan Petkov, Lovisa Franzén, Jessica Bunz, Christine Moge, Henrik Everberg, Petter Brodin, Alvaro Martinez Barrio, Simon Fredriksson

## Abstract

Cellular functions depend on dynamic interactions of proteins and their spatial organisation. While transcriptomic and proteomic methods of molecular parts-lists have enabled single-cell profiling based on abundance, scalable technologies allowing high-resolution measurements of protein organization and interactions at scale are lacking. Here we present the Proximity Network Assay (PNA), a DNA-based method for constructing three-dimensional nanoscale maps of 155 plasma membrane proteins in single cells without the use of optics. PNA employs barcoded antibodies and *in situ* rolling circle amplification to generate >40,000 spatial nodes per cell, which are linked through proximity-dependent gap-fill ligation and decoded by DNA sequencing forming single cell Proximity Networks. PNA captures abundance, self-clustering, and ∼12,000 pairwise colocalization relationships per single-cell, validating established protein interactions. This new modality provides a framework to uncover novel spatial biomarkers, reveal functional mechanisms, and advance translational studies in immunology, oncology, and cell therapy.

## Introduction

Proteins function in dynamic clusters with other proteins to carry out cellular activities, including migration, adhesion, and signal transduction^1–4^. These assemblies encode regulatory states and dynamic interactions that cannot be inferred from abundance alone. Approximately 60% of all known drug targets reside in the plasma membrane and are functionally dependent on the nanoscale organisation and clustering of membrane-associated proteins^5,6^. However, measuring spatial protein organization at high resolution and throughput for single-cells remains a fundamental challenge.

Despite the availability of high-throughput omics technologies for transcript and protein quantification at the single-cell level, scalable tools to measure spatial protein interactions with nanoscale resolution and single-cell granularity are lacking. Imaging-based methods such as fluorescence microscopy and imaging mass cytometry are limited by optical resolution and multiplexing constraints due to their reliance on light. Proximity-based methods like *in situ* Proximity Ligation Assay (PLA), Prox-seq, and Förster Resonance Energy Transfer (FRET) achieve spatial resolution down to 10–40 nm but are inherently limited to pairwise proximity interactions and does not make networks or multiprotein domains^7,8^. In contrast, mass spectrometry-based approaches such as co-immunoprecipitation (co-IP) or proximity labeling techniques like BioID (proximity-dependent biotin identification) and APEX (engineered ascorbate peroxidase) provide broader, unbiased discovery capabilities^9^, but require large, homogeneous cell populations (typically millions of cells), lack spatial resolution, and are incompatible with profiling of heterogeneous clinical samples. Collectively, these limitations have left a critical gap in our ability to study protein organization and interactions at the scale, resolution, and throughput needed for single-cell translational research applications.

We previously introduced Molecular Pixelation (MPX); a DNA-based, optics-free spatial proteomics method that segments the cell surface of individual cells into zones using barcoded antibody-oligonucleotide conjugates and spatially anchored concatemeric UMIs at ∼280 nm resolution^10^. MPX, part of the emerging field of DNA microscopy^11–16^, alleviated the limitations of fluorescence multiplexing by encoding spatial proximity in DNA sequence space.

Here, we present the Proximity Network Assay (PNA), a next-generation sequencing-based platform for reconstructing three-dimensional spatial Proximity Networks of proteins in single cells. PNA builds on the core principle of proximity-based assays such as PLA and Proximity Extension Assay (PEA)^17–19^, but moves beyond pairwise designs by introducing multiple spatial contact points per protein target. From each barcode, attached to an antibody, an *in situ* rolling circle amplification (RCA) event is initiated, generating many identical barcode copies that serve as spatial nodes. These nodes are linked to neighboring RCA products via hybridisation of a linker oligonucleotide, followed by gap-fill ligation to generate united barcode pairs. These DNA constructs, representing spatial proximity events, are read out by Next-Generation Sequencing (NGS) and computationally reconstructed into Proximity Networks; graph-based representations of single-cell and single molecule nanoscale networks.

We demonstrate the performance of PNA by profiling 155 antibody-targeted proteins with the Pixelgen Proxiome kit, generating ∼40,000 spatial nodes per cell at single molecule resolution. PNA avoids the limitations of microscopy readout and integrates with standard NGS workflows. The resulting data structure supports analysis of protein abundance, spatial self-clustering, and colocalization, allowing for a systems-level view of protein organisation.

We apply PNA to a diverse set of biological systems including peripheral blood mononuclear cells (PBMCs), hematologic cancer cell lines, CAR-T cell models, and autoimmune disease samples, demonstrating its ability to resolve protein interactions, spatial rearrangements, and cell type specific network topologies. By enabling nanoscale spatial proximity mapping of thousands of single cells in clinical and experimental samples, PNA opens a new dimension for studying protein networks at scale, with applications in systems immunology, drug target discovery, biomarker discovery, and translational research.

## Results

### Proximity Network Assay

PNA operates in suspension, using standard molecular biology workflows (**Figure 1A**). PFA fixed cells are bound by barcoded monoclonal antibodies (Ab-BC) targeting surface proteins. Each Ab-BC exists in two versions (type-1 and type-2) with distinct oligonucleotide sequences to avoid downstream self-interaction events. These oligonucleotides carry a Protein ID (PID) and a unique molecular identifier (UMI), and are hybridized by type-1 or type-2 gap-fill ligation padlock probes to which the PID and UMIs are copied upon circularization. The circularized padlock is amplified by a short *in situ* RCA, generating a localized cluster of repeated barcode copies anchored at the site of protein binding (**Figure 1A**).

**Figure 1.**
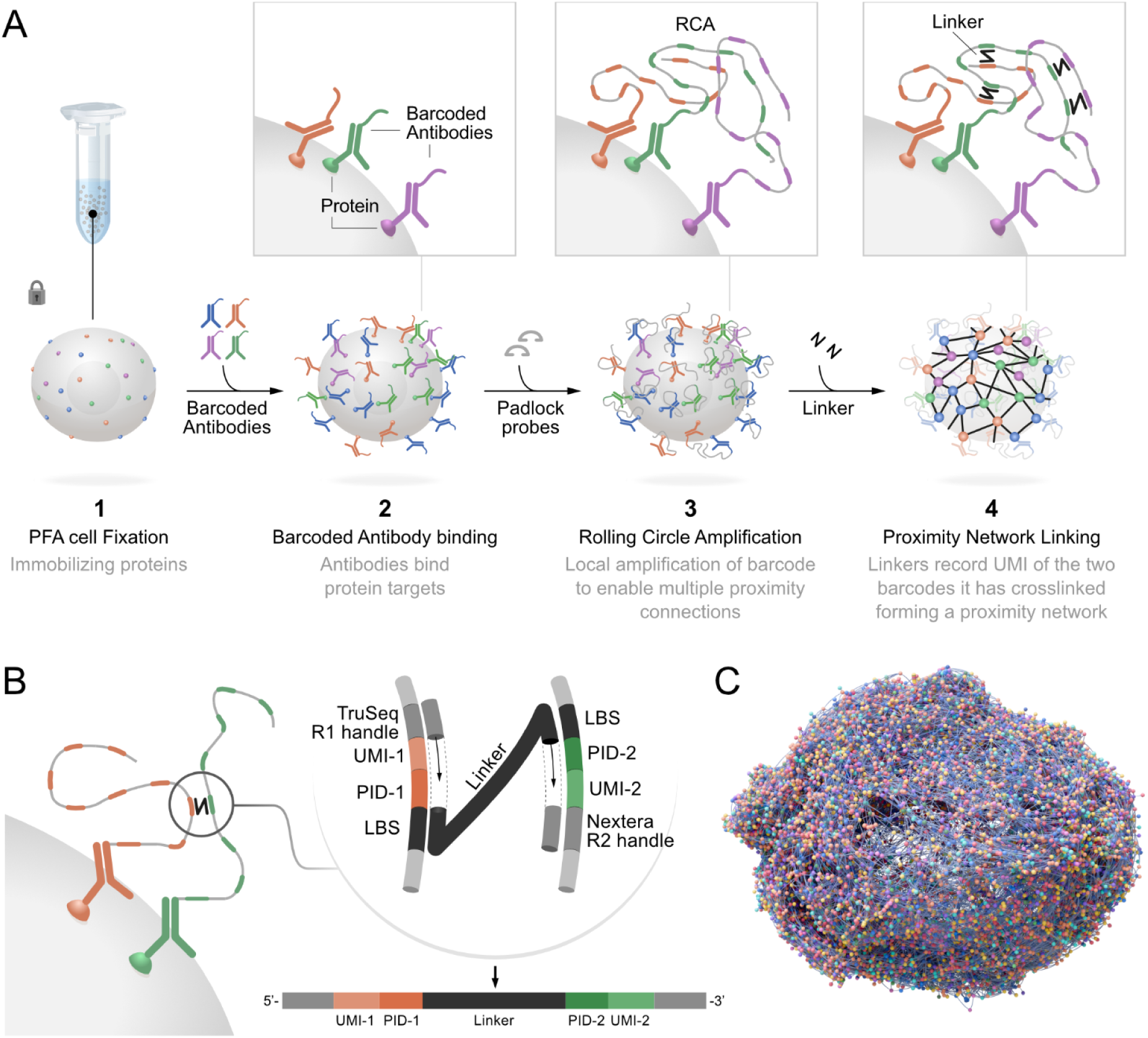
The Proximity Network Assay connects surface proteins into a cell-wide network. (A) Proximity Network process steps. 1. Cells are fixed using PFA, 2. Ab-BCs are bound to PFA fixed cells in suspension. 3. Padlock probes are hybridized, followed by gap-fill ligation copying the UMI and Protein ID (PID) of the Ab-BC into the circularized padlock. RCA then makes multiple copies of the UMI and PID. 4. Linker oligonucleotides are hybridized to pairs of proximal RCA products. (B) Overview of linker amplicon construct: Gap-fill ligation incorporates UMI and PID into the target amplicon from two RCA products of type 1 (orange) or type 2 (green), respectively. The amplicon is subsequently PCR:ed and sequenced. (C) 3D reconstruction of the Proximity Network of a single cell. Each node corresponds to a protein, and each connection a linker molecule.

To detect spatial proximity events, linker oligonucleotides are introduced that hybridize to two RCA products in close proximity to each other; specifically, one type-1 and one type-2 product. Because the linker is designed to bind exactly one type-1 and one type-2 binding site, it cannot hybridize to binding sites within a single RCA product preventing single molecule self-interaction events. A subsequent gap-fill ligation step incorporates both barcodes (PID and UMI) from each RCA product onto the hybridized linker oligonucleotide along with primer sites for PCR to enable amplification (**Figure 1B**). Each generated DNA fragment therefore represents a proximity link between two proteins.

Unlike methods that require physical isolation of single cells (e.g. droplet barcoding or imaging), PNA does not require compartmentalization. Instead, each cell is represented by a unique combination of tens of thousands of interconnected UMIs making up its Proximity Network. As a result, over a thousand cells can be assayed together in one tube, and later deconvoluted into single-cell Proximity Networks representing the spatial arrangement of the 155 targeted proteins.

The resulting product is a densely connected network for each single cell in 3D (**Figure 1C and Supplementary Video 1**) composed of uniquely positioned protein molecules, each spatially connected to a median of 5 other protein molecules on PBMCs (**Supplementary Figure 1A**). To demonstrate the cohesion of the network, the PNA protocol was conducted using fluorescently labeled nucleotides during the *in situ* RCA reactions, and visualized by microscopy **(Supplementary Figure 2**). The PNA reaction products localized specifically to the cell membrane of each single cell and demonstrated even coverage over the entire cell surface.

Since the generated Proximity Networks are spatially resolved, PNA enables not only analysis of protein abundance, but also analysis of spatial patterns across cells, such as spatial self-clustering of each protein and colocalization between different proteins. Due to the intrinsically high multiplex, the assay uniquely profiles the spatial relationships of thousands of protein pair combinations on single cells, simultaneously. The spatial resolution of the method is estimated to 50 nm by dividing the surface area of a lymphocyte, 92 um^2, by the approximately 40,000 unique spatial positions of each cell network^20^ (**Supplementary Figure 1B**). The absolute resolution is dependent on RCA product size and degree of RCA product compaction, reported below 100 nm by electron microscopy^21^. Larger cells do have more spatial positions and require more sequencing reads such as Raji-cells.

We developed a Proximity Score based on the join count statistic^22^ to summarize the level of self clustering or colocalization of proteins. Comparison of the observed join counts with a null distribution from random permutations uniquely adjusts for abundance-driven colocalization and allows for statistical interpretation of significance of each observation. This ability to study colocalization and clustering in isolation from abundance is challenging in many commonly used methods for protein interaction analyses such as *in situ* PLA, fluorescence-based colocalization, proximity labeling, or FRET. Abundance level data in PNA can be analyzed in a similar fashion as other single-cell abundance based methods.

It is important to note that due to the monoclonal nature of the antibodies and the specific type-1 and type-2 Ab-BCs formulation described above, homo-cis (same protein, same molecule) data is not generated as only one antibody-barcode can occupy a single target protein molecule and self-interactions events are not generated. Instead, the Proximity Score captures hetero-trans (different proteins) and homo-trans (same protein, different molecules) interactions corresponding to colocalization and clustering quantified in the generated data.

PNA offers several advantages over the previously introduced MPX. PNA shows improved stability in spatial scores, enabling robust detection of spatial signatures even with very few cells (**Supplementary Figure 3A**). Furthermore, PNA demonstrates increased sensitivity for detecting known interactions (**Supplementary Figure 3B-C**), exhibits superior network metrics (**Supplementary Figure 3D**), and, perhaps most notably, PNA marks a substantial improvement in spatial sampling. Specifically, the total number of sampled spatial locations is an order of magnitude higher than in MPX, with single molecule granularity (**Supplementary Figure 3E-F**).

### Proximity Scores detect known protein interactions

To demonstrate spatial analysis by PNA, we assayed 1,017 cells of the Burkitt’s lymphoma cell line Raji, a cell line of B cell origin. The Proximity Score used for spatial characterization is formulated as a log_2_ ratio between the observed number of join counts and the expected value under a random spatial model, conditional on the observed abundances of the two markers. Consequently, a score of 1 indicates that the two markers occur in proximity of each other twice as often as expected by chance, while a score of 2 corresponds to a 4-fold enrichment. Conversely, a score of −1 indicates half the expected frequency, and −2 corresponds to a 4-fold depletion. Thus, a positive score is indicative of colocalization, whereas negative scores reflect spatial segregation between markers. The expected join count was estimated from a spatial null model generated using 100 random permutations, after which Proximity Scores showed negligible variation with additional permutations (**Supplementary Figure 4A-B**).

The Proximity Scores of Raji cells revealed an intricate landscape of colocalizing and segregating proteins, attesting the existence of multiple protein complexes and microdomains at the cell surface (**Figure 2A**). These include well-known interacting protein pairs such as MHC I (HLA-ABC/B2M), LFA-1 (CD11a/CD18), the B cell receptor (BCR) (CD79a/IgM), and VLA-4 (CD49D/CD29). We could also identify colocalization patterns distinctive of known microdomains including CD20 colocalizing in tetraspanin-rich microdomains (CD37/CD53/CD81/CD82)^23^, and GPI-coupled receptors (CD48/CD52/CD55/CD58/CD59) associated with lipid rafts^24^. As expected, high Proximity Scores were found for protein isoforms detected by two probes targeting the same protein but at different epitopes, such as HLA-DR with HLA-DR-DP-DQ, and CD45 with CD45RA and CD45RB.

**Figure 2.**
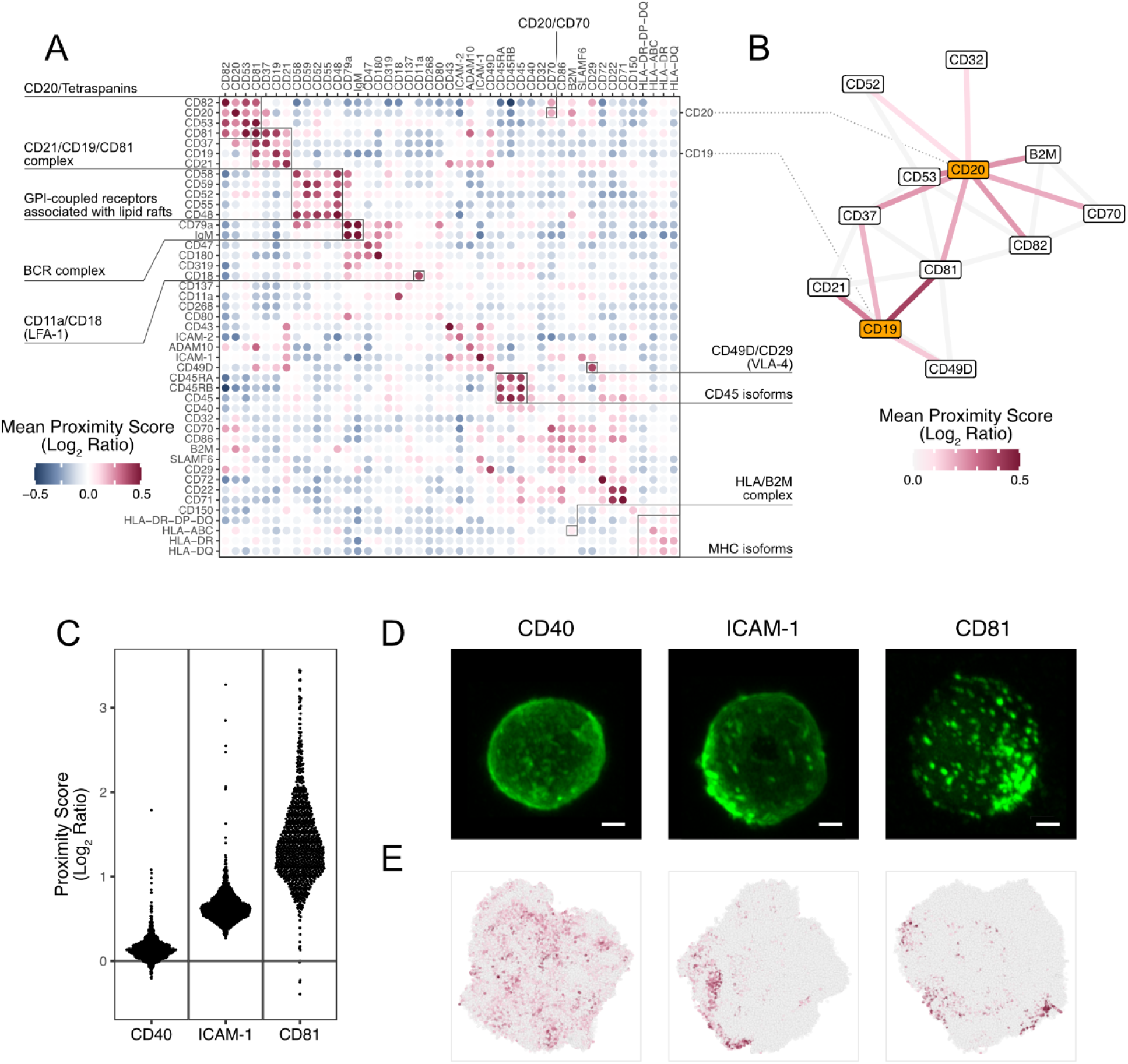
Spatial organization of Raji cell surface proteins. (A) Heatmap of mean Proximity Scores (log2 ratio) of Raji cells showing protein colocalization and self-clustering of abundant proteins. Black frames highlight examples of known protein-protein interactions and previously described multi-protein domains. (B) The interaction network of CD19 and CD20, composed of proteins showing mean Proximity Scores above 0.1 with CD19 or CD20. Gray lines indicate mean Proximity Scores above 0.1 between other members of this interaction network. (C) Proximity Scores across Raji cells for the markers CD40, ICAM-1, and CD81. Each point corresponds to the score of a single cell. (D) Representative immunofluorescence (IF) microscopy images of single cells, showing the clustering behavior of CD40, ICAM-1 and CD81. Scale bars: 2 μm. (E) Representative 3D visualizations of PNA cell graphs of single cells showing the distribution of CD40, ICAM-1, and CD81. A darker dot color indicates a high local density of the protein in the graph, here quantified as the Getis-Ord statistic.

We then focused on two key proteins of B cell function, and generated a protein interaction network centered on CD19 and CD20 (**Figure 2B**). These two networks were connected since CD19 and CD20 both showed high colocalization with tetraspanins CD37 and CD81. We could detect the recently described interaction between CD20 and CD70^25^, as well as the B cell co-receptor complex formed by CD19 associating with CD81 and CD21, which plays an important role in regulating and enhancing signalling through the BCR^26,27^. Interestingly, besides this interaction network, CD21 was also associated with a distinct multiprotein domain rich in adhesion molecules including ICAM-1 and ICAM-2.

In addition, Proximity Scores were frequently elevated for proteins with themselves, reflecting spatial clustering, in agreement with previous reports in literature^28^. Proximity Scores generated by PNA of self-clustering proteins show concordance with immunofluorescence (IF) microscopy images, here exemplified by three proteins with low to high level of clustering (**Figure 2C-E**, **Supplementary Figure 5A-C**). CD40 displayed an even distribution across the cell surface, while ICAM-1 and CD81 showed increasingly higher Proximity Score as well as clustering in IF (**Figure 2C-E**) in concordance with literature on CD81^29^. While some cells exhibit unusually high or low Proximity Scores, these outliers are not associated with differences in graph size or connectivity across QC metrics (Supplementary Fig. 5D).

### Deep phenotyping of heterogenous immune cell populations

The 155-plex panel of the Proxiome kit was developed to cover the most important surface markers to characterize the function and identity of key human immune cell types. To demonstrate the breadth of the panel and the specificity of the included antibodies, we performed a deep annotation of cell types in PBMCs (**Figure 3A-B**). Utilizing a hierarchical annotation strategy (**Supplementary Figure 6**) where cells were divided into gates of increasing granularity to a total of 34 cell type identities, out of which we were able to identify cells of 30 identities, including subtypes of all major PBMC lineages; T cells, B cells, natural killer cells (NKs), dendritic cells (DCs), monocytes, but also small populations of common PBMC contaminant cell types; platelets (CD41^+^, CD45^-^) and basophil granulocytes (CD66b^+^, CD193^+^). **Figure 3A** shows a UMAP based on protein abundance where cell types identified by gating form separate clusters according to expected patterns, exemplified by that various memory compartments of CD4 and CD8 T cells, respectively, clustering together, but distinct from their respective naive CD4/CD8 T cell population.

**Figure 3.**
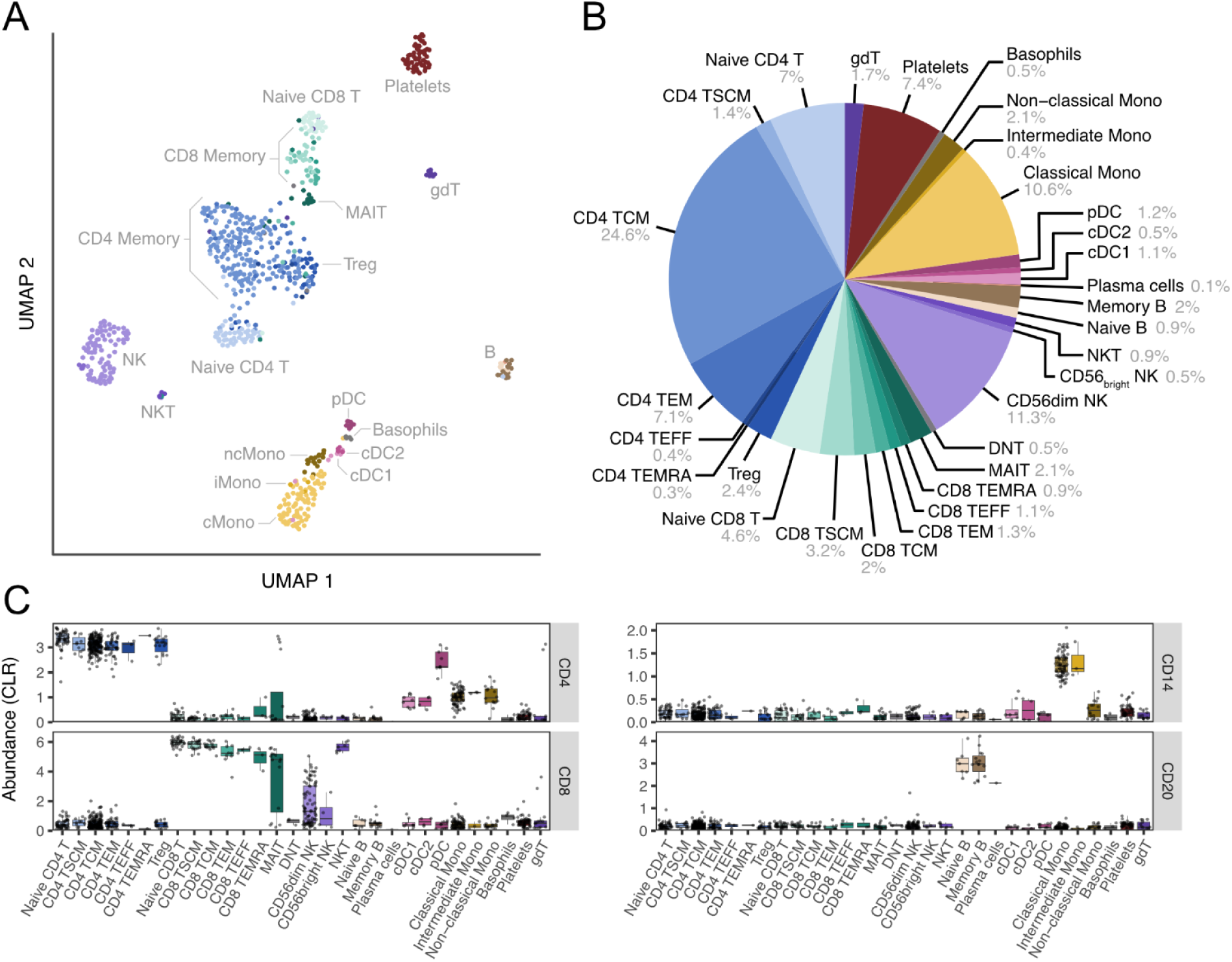
High-dimensional cell type annotation of PBMC samples. (A) UMAP of cell annotation in PBMC. (B) Proportions of found cell types based on abundance of canonical lineage markers. (C) Boxplot of the abundance of markers CD4, CD8, CD14, and CD20 in the various cell types identified. Abbreviations: TSCM - stem cell memory T cells; TCM - central memory T cells; TEM - effector memory T cells; TEFF - effector T cells; TEMRA - terminally differentiated effector memory T cells; Treg - regulatory T cells; MAIT - mucosal-associated invariant T cells; DNT - double-negative T cells; NK - Natural Killer cells; NKT - Natural Killer T cells; cDC - conventional dendritic cells; pDC - plasmacytoid dendritic cells; Mono - Monocytes; gdT - gamma delta T cells.

Identified cell types show expected abundance patterns (**Figure 3C** and **Supplementary Figure 7**), with lineage markers such as CD14 and CD20 distinguishing monocytes and B cells, respectively, and examples of population exclusive expression of T cell receptor (TCR) variants TCRgd, TCRVd2, and TCRVg9 in the gamma delta T cell (gdT) population, which is also the only T cell population that does not express TCRab. To validate the cell annotation, we performed flow cytometry in addition to PNA to quantify a selection of cell populations using an orthogonal method across resting PBMC samples from three different donors. The result showed high concordance between PNA and flow cytometry, exemplified by T cells (CD45^+^, CD3^+^, PNA: 58.2% vs Flow: 59.4%) (**Supplementary Figure 8A**), as well as the CD4^+^ (PNA: 57.8% vs Flow: 61.1%) and CD8^+^ (PNA: 32.8% vs Flow: 28.2%) subsets of T cells (**Supplementary Figure 8B**). The overall correlation of estimated population sizes was very high, with a Pearson’s r = 0.99 between cell population sizes identified by PNA and Flow cytometry (**Supplementary Figure 8C**). These results confirm the robustness and accuracy of PNA-based cell type annotation in our dataset.

### CAR-T cell Proximity Networks

To evaluate the capacity of PNA for resolving dynamic remodeling of engineered immune cells, we profiled CD19-targeting CAR-T cells before and after coculture with CD19⁺ Raji tumor cells (**Figure 4**). CAR-T cells were analyzed either in isolation or following coculture at an effector-to-target (E:T) ratio of 1:1 for 4 or 24 hours (**Figure 4A**). Each sample was processed using the 155-plex PNA panel supplemented with a CAR-specific anti-FMC63 probe.

**Figure 4.**
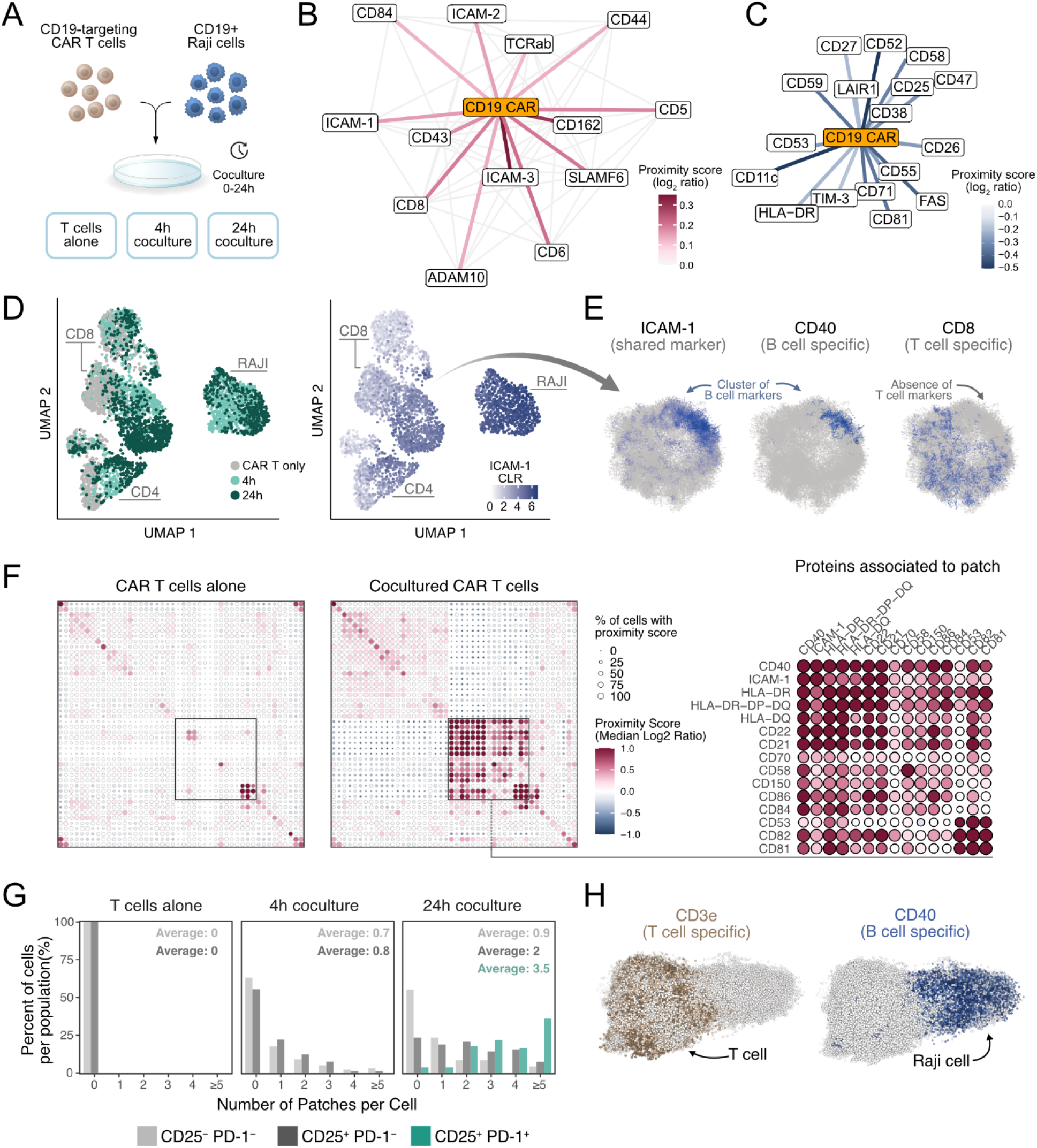
The Proximity Network Assay resolves CAR-T cell surface remodeling, trogocytosis, and tumor cell conjugation. (A) Workflow for functional characterization of CAR-T cells. CAR-T cells were either kept separate, or were mixed with CD19-positive Raji cells at an 1:1 E:T ratio. Cells were cocultured for either 4h or 24h, after which they were retrieved and fixed using 1 % PFA. Each sample was split into two duplicates and processed with the Proximity Network Assay according to the DP001 CAR Barcoded Antibody spike-in Proxiome kit (v1.01) manual. (B) Proteins displaying positive colocalization (average score ≥0.1) with the CD19 CAR were plotted in a network. Positive colocalization scores for protein pairs other than the CD19 CAR have been grayed out for clarity. (C) Proteins displaying negative colocalization (average score ≤-0.1) with the CD19 CAR were plotted in a network. (D) Left: High-dimensional clustering based on protein abundance data for CAR-T cells alone, 4h cocultured cells, and 24h cocultured cells efficiently separating CD4, CD8 and Raji cells. Right: Cocultured T cells displayed increased levels of ICAM-1. (E) 3D visualization of a single CD8 T cell from the 24h cocultured sample. The cell displays a patchy ICAM-1 distribution where the patch colocalized with B cell markers like CD40, while anti-colocalizing with T cell marker CD8. A darker dot color indicates a higher local density of the protein. (F) Colocalization heatmaps of abundant proteins in unstimulated (left) and 24h cocultured (right) CAR T cells, displaying the acquisition of a protein domain following coculture. (G) Quantification of tumor patches on the surface of CD8^+^ non-activated (CD25^-^, PD-1^-^), activated (CD25^+^, PD-1^-^) and exhausted (CD25^+^, PD-1^+^) T cells. (H) An example of a CD8 T: Raji cell conjugate. The T cell (left) displays expression of T cell specific marker CD3e, while the Raji cell (right) expresses B cell marker CD40. The nodes are colored by density of the markers, measured by the Getis-Ord statistics. Only positive nodes are colored.

To characterize CAR-associated interaction landscapes, we computed Proximity Scores between the CD19 CAR and all other panel markers, generating a network of positively colocalized proteins and one of segregating proteins (**Figure 4B-C** and **Supplementary Figure 9A**). The CAR receptor exhibited spatial proximity with canonical T cell signaling and adhesion proteins, including TCRαβ, CD5 and CD6, and multiple ICAM family members. In contrast, the CAR is segregated from lipid raft-associated markers such as CD52, CD55 and tetraspanins CD53, and CD81. These results indicate that the synthetic CD19 CAR receptor is not randomly distributed but integrates into endogenous TCR-associated signaling domains while separating from raft-like compartments, partly mirroring native TCR topology.

Progressive remodeling of the T cell surface proteome was observed during coculture. UMAP clustering of samples based on marker abundance showed clear shifts between monoculture, 4-hour, and 24-hour conditions, with pronounced upregulation of immune activation markers, and downregulation of key activating receptors including the CAR, the TCR and CD28 (**Supplementary Figure 9B)**. Notably, ICAM-1 expression increased substantially following coculture (**Figure 4D**). However, 3D spatial rendering revealed that ICAM-1 enrichment occurred in discrete patches rather than uniformly across the cell surface (**Figure 4E)**.

By measuring spatial scores across the whole panel, we could reveal the formation of a highly colocalized protein domain at the surface of cocultured CAR T cells (**Figure 4F and Supplementary Figure 10**). Besides ICAM-1, this domain contained numerous B cell–specific markers such as CD40 and CD21, while segregating from T cell markers like CD8 (**Figure 4E-F, Supplementary Figure 9C**). These findings are consistent with trogocytosis, an important, yet poorly understood mechanism of CAR T cell regulation, whereby membrane fragments are transferred between target and effector cells during cell interaction^30,31^. Interestingly, we observed similar reorganization patterns at the surface of T cells following PHA-stimulation of PBMCs. These activated T cells displayed acquisition of B cell patches, indicating interaction and trogocytosis between cells also in a non-cytotoxic setting (**Supplementary Figure 11A-D**).

To explore the relationship between patch acquisition and CAR T cell state, we stratified CD8⁺ cells by canonical activation and exhaustion markers. Cells classified as non-activated (CD25⁻ PD-1⁻) exhibited few or no patches, while activated cells (CD25⁺ PD-1⁻) carried an intermediate patch burden. Exhausted cells (CD25⁺ PD-1⁺) displayed the highest frequency and number of tumor patches, with a ∼5-fold increase in patch count relative to baseline (**Figure 4G)**. These results suggest that PNA enables functional stratification of immune effector cells based on spatial acquisition of tumor-derived proteins, with the content of these patches providing clues to the history of cell-cell interactions.

In addition to detecting trogocytosis, PNA enabled identification of direct T cell-tumor cell conjugates. Proximity Networks of conjugated cell pairs revealed clear localization of T cell-specific markers (e.g. CD3e) on one side and B cell markers (e.g. CD40) on the opposite pole (**Figure 4H**). This pattern was distinct from trogocytic patching and consistent with immune synapse formation. Such conjugates can be distinguished by PNA from trogocytic cells, providing a high-resolution approach for identifying physical immune engagements and mapping the molecular topology of intercellular contact interfaces.

Together, these results demonstrate that PNA captures the nanoscale organization and dynamic remodeling of the CAR-T cell surface during tumor interaction, including co-receptor colocalization, ligand acquisition via trogocytosis, and synapse formation.

### Systemic lupus erythematosus (SLE) study

To assess whether protein Proximity Networks can serve as a novel class of spatial biomarkers, we profiled B cells from patients with systemic lupus erythematosus (SLE, n = 2) and healthy controls (Controls, n = 2) (**Figure 5A-B**) (Figure 5A-B). SLE is widely understood as a disorder of impaired B cell tolerance and dysregulated B-cell activation, making this population particularly relevant for biomarker discovery. We therefore examined whether disease status is reflected not only in protein abundance but also in the protein organization of B cells through their Proximity Networks (**Figure 5C**), thereby evaluating receptor organization as a potential biomarker dimension in autoimmune disease.

**Figure 5.**
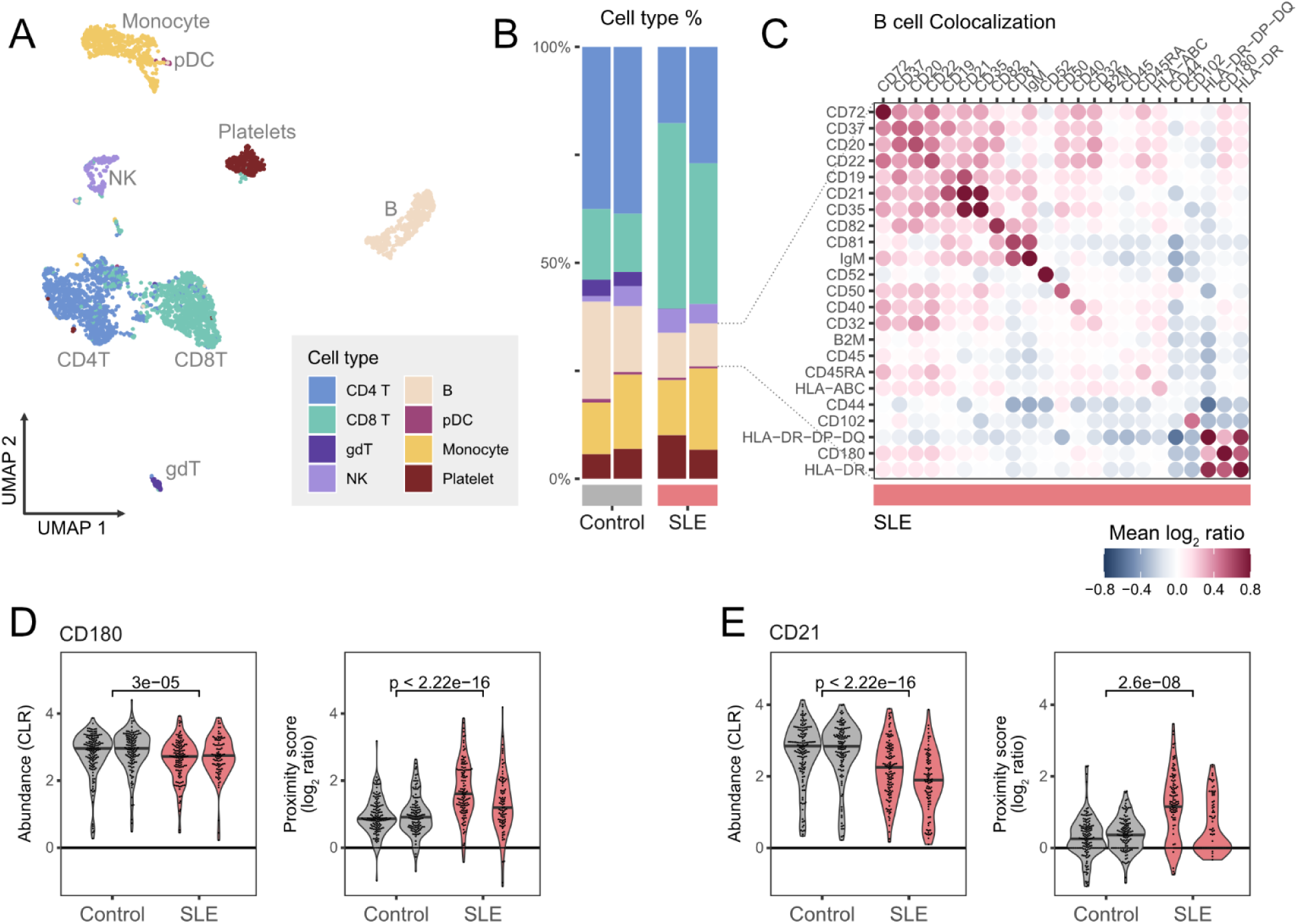
SLE B cells display increased clustering of CD180 and CD21. Protein abundance and spatial organization was assessed for SLE compared to healthy B-cells from two PBMC samples of each type. (A) UMAP of annotated PBMC cells of 2 healthy control samples and 2 SLE samples. (B) Bar plots of the cell composition of the samples. (C) Proximity Score heatmap of the 30 markers with highest mean abundance (CLR) among SLE B cells. (D) Violin plots of abundance (left) and Proximity Score (right) of CD180. (E) Violin plots of abundance (left) and Proximity Score (right) of CD21.

We observed reduced surface abundance of CD180 (RP105) and CD21 (complement receptor 2, CR2) in SLE B cells compared to controls (CD180: Wilcoxon p = 3 × 10^-5^; CD21: Wilcoxon p < 2.22 × 10^-16^) (**Figure 5D–E**). CD21 is a core component of the B cell co-receptor complex, where it enhances signalling through the B cell receptor (BCR)^27^. Reduced CD21 expression on B-cells in SLE-patients has previously been reported and associated with high disease activity^32^. CD180 was previously reported as associated with higher SLE disease activity^33^. Low B cell abundance of CD180, a Toll-like receptor that counter-regulates type I IFN, has been linked to higher IFNα signalling in lupus-prone mice^34^.

Despite their reduced abundance, both CD180 and CD21 exhibited significantly higher Proximity Scores in SLE compared to controls (CD180: Wilcoxon p < 2.22 × 10^-16^; CD21: Wilcoxon p = 1.6 × 10^-8^), indicating increased spatial clustering within restricted membrane regions (Figure 5D–E). Notably, the abundance reduction was modest for CD180, suggesting that altered spatial organization is not simply a secondary effect of differential expression. These findings demonstrate that proximity-based metrics can capture changes in membrane topology that are not apparent from abundance measurements alone.

Although the functional consequences of this polarized receptor distribution remain to be determined, altered spatial organization of CD21 and CD180 may influence local signal integration at the B cell surface. For instance, clustering of CD21 could serve to focus and amplify local signalling from complement-decorated immune complexes in B cells and could be a secondary consequence of increased immune complexes in circulation forcing CD21 molecules into proximity. Alternatively, it may represent a compensatory mechanism by B cells to remove CD21 from active signalling hubs and dampen chronic activation. Similar mechanisms could apply to CD180, or such receptor clustering could instead represent a disease-driving phenotype where limiting receptor polarization might alleviate disease progression and patient symptoms.

The PNA platform offers a promising approach to longitudinally monitor and profile receptor topology in SLE patients, track changes during disease flares, and assess whether biological therapies can restore a more physiological, symmetric distribution of CD21, CD180, and related receptors as a novel therapeutic modality. Further investigation will be required to understand the consequences of such polarized localization.

## Discussion

The functional state of a cell is defined not only by its transcriptomic or proteomic composition, but critically by the spatial organisation of its proteins through clustering, segregation, and interactions. This organization governs essential cellular processes such as signal transduction, immune activation, and cell-cell communication. Yet, despite its biological significance, spatial protein organisation remains largely unexplored at the single-cell, omics-scale in heterogeneous clinical samples.

Existing tools for protein interaction studies either lack throughput or resolution, or require bulk input from homogeneous cell populations. For instance, proximity labeling (PL) techniques such as BioID and APEX, read out by mass spectrometry, have enabled discovery of thousands of interaction partners in a bulk format, but require tens of millions of cells and achieve limited resolution (∼200-300 nm)^35,36^, These approaches are unsuitable for studying heterogeneous clinical samples, where each cell type requires a separate analysis to enable biomarker discovery and translational research.

The Proximity Network Assay addresses this unmet need by enabling high-throughput, single-cell, nanoscale single molecule protein networks in a DNA-sequencing-based readout. The presented implementation of PNA for cell surface analysis captures the proximity profile of 155 antibody-targeted proteins, with demonstrated applications in human immune cells, cell lines, and CAR-T cells. This is achieved without the need for single cell compartmentalization as the method generates an interconnected spatial network of barcodes for each assayed cell.

PNA extends the conceptual framework of DNA-based proximity methods such as PLA and PEA^16,17^, which rely on pairwise proximity detection via sequence generation. Unlike these pairwise assays, PNA constructs comprehensive networks of protein proximities, enabling analysis of abundance, spatial self-clustering of one protein species, and colocalization of protein pairs. This generalisation from binary interactions to spatial network topology distinguishes PNA from prior proximity technologies including Prox-seq^8^. It also represents an advancement over the Molecular Pixelation (MPX) platfor^10^, which demonstrated optics-free spatial reconstruction but with zone-based lower spatial resolution compared to PNA single molecule resolution. As all antibody based methods, PNA is limited by antibody availability and potential steric hindrance of epitopes by dynamics of post-translational modifications.

PNA is validated by its ability to detect well-characterised molecular complexes, such as HLA-ABC/β2-microglobulin, CD19/CD81, IgM/CD79a, integrin complexes, and more. Spatial proximity metrics may serve as interaction-based biomarkers, including for response prediction to immune checkpoint inhibitors^37^, or for investigating CAR-T cell phenotypes^38^. However, as with all proximity-based approaches, a high Proximity Score does not confirm direct physical interaction. It may reflect nanoclustering, shared membrane microdomains, or subcellular colocalization^39,40^. This distinction is important when interpreting interaction candidates. Nevertheless, the reproducible detection of known complexes and consistent clustering patterns supports the utility of PNA in hypothesis generation, mechanistic inference, clinical biomarker discovery, and support development of therapy-induced spatial remodelling of surface proteins^41^. The identification of novel protein proximities that are prevalent on cancer cells and aberrant immune cells in comparison to healthy cells is of particular interest, as these may constitute novel drug targets using bispecific antibodies and CAR-T cells potentially increasing the number of druggable cell surface targets.

The graph-native structure of PNA data opens new analytical possibilities suited for machine learning, including graph convolutional networks (GCNs) and graph attention models^42^. These algorithms can identify latent structure, classify spatial phenotypes, and predict cell state transitions or therapeutic outcomes from network topology.

Looking forward, the potential of PNA is considerable. Preliminary findings show that Proximity Networks can also be generated within cells, enabling nanoscale localization of intracellular proteins in signaling pathways, immune synapse protein interactions between cells, and single cell transcriptomics using RNA probes (work in progress).

The Proximity Network Assay has the potential to impact our understanding of single cell protein organization and interactions beyond what’s achievable through traditional microscopy which is limited in resolution, multiplexing, and throughput. PNA requires approximately 300,000 sequencing reads per cell and is compatible with existing NGS platforms based on sequencing-by-synthesis (SBS) and also using higher throughput nanopore sequencing sequencing-by-expansion (SBX)^43^ making the entire work flow completely optics-free (**Supplementary Figure 12A-E**). As sequencing technologies continue to advance, the output of PNA data will increase further, making analyses of larger sample areas such as tissue feasible.

Current single cell research is mostly based on parts-list abundance measurements of transcripts and proteins per cell. Single-molecule Proximity Networks enabled by PNA adds a new analysis modality, to gain novel insights into the molecular mechanisms of cell function and for biomarker discovery.

## Acknowledgements

Author Contributions Statement

FK and SF conceived of the Proximity Network Assay. FK, SF, MiS, CG, TK, DT, HvO, LL, MK, JB, CM, MaS, HvO, PB, SG, VT, RF, EN, BW, LX developed the method and/or performed experiments. FK, AMB, JD, FdT, PT, LL, MK, developed the data analysis algorithms and the pipeline. FK, MK, VvH, AMB, PB, HvO, DT, SP, LF, RF, PB performed in depth analyses and interpretation of biological results. FK, MK, HvO, VvH, PB, and SF wrote the manuscript. All the authors have read and approved the manuscript before submission.

## Competing interests

All authors (except RF, EN, BW, and LX) are employees or advisors to Pixelgen Technologies AB commercializing products based on Proximity Networks. They are also shareholders, advisors, or stock option holders of the company. Proximity Networks and Proxiome are trademarks of Pixelgen Technologies AB. BW and LX are employees of Roche Diagnostics.

## Ethics declaration

All samples acquired from the Karolinska Hospital with informed consent and withheld sample identity or other medical information.

## Methods

### PBMC extraction from buffy coats

Buffy coats derived from healthy, anonymous donors, were acquired from Komponentlab, Karolinska hospital. The blood was diluted and the PBMCs were isolated through density gradient centrifugation using Lymphoprep (Stemcell technologies). The platelet fraction was reduced by 3 repeated low-speed centrifugation steps, after which remaining red blood cells were lysed using eBioscience RBC lysis buffer (Invitrogen). Resting cells were fixed using 1% paraformaldehyde (PFA) and frozen for long-term storage.

### Cell culture

Raji cells were cultured in RPMI media supplemented with 10% fetal bovine serum and 1% Penicillin-Streptomycin and were split every 48 hours. Cell cultures were regularly confirmed negative for mycoplasma contamination.

### CD19 CAR-T cell - tumor cell coculture

CD19-targeting CAR-T cells (Promab) were thawed and washed, after which they were fixed using 1% PFA, or cocultured with Raji cells at an E:T ratio of 1:1 for 4 or 24 hours in U-bottom 96-well plates. After the indicated time, cocultured cells were harvested and fixed using 1% PFA, and frozen and stored before running PNA.

### Microscopy of PNA products

Millicell® EZ Slides were coated with 0.1% (w/v) poly-L-lysine for 5 minutes, then air-dried at room temperature. PBMCs were fixed and blocked according to the PNA protocol. The RCA reaction was spiked with 1ul of 1mM Fluorescin-dUTP. After RCA, cells were washed twice, and subsequently stained with AF647-conjugated Wheat Germ Agglutinin (WGA). The suspension was added to a coated EZ Slide well and incubated overnight at 4 °C to allow cells to settle. The next day, slides were mounted with SlowFade™ Glass Soft-set Antifade Mountant containing Dapi. Slides were imaged in 3D on a Zeiss LSM 980 using the Airyscanning module and a 63X (1.4) oil objective. Analysis was performed using ImageJ. Images were corrected for spectral drift and brightness and contrast adjusted. 3D stack were compiled, and intensity profiles were measured for single optical planes for the WGA and PNA-product channels.

### Comparison of PNA and flow cytometry for cell type identification

PBMC cells from 3 different donors were fixed using 1% PFA, washed and analyzed using PNA, or stained for flow cytometry using CD45 Pe, CD3 PeCy7 and CD56 APC, or CD45 Pe and CD8 AF488, or CD45 Pe and CD4 Fitc, or CD45 Pe and CD19 AF700, or CD45 Pe and CD14 APC, or CD4 Pe, CD3 PeCy7 and CD8 AF488. Cells were processed on a CytoFLEX (Beckman Coulter) and analyzed using FlowJo. Cells were gated for: (CD14^+^, CD45^+^), (CD19^+^, CD45^+^), (CD3^+^, CD45^+^), (CD56^+^, CD45^+^), (CD8^+^, CD3^+^), (CD4^+^, CD3^+^), (CD3^+^, CD4^+^, CD8^-^), (CD3^+^, CD8^+^, CD4^-.^).

### Microscopy of protein clustering in Raji cells

Raji cells were fixed using 1% PFA, washed and analyzed using PNA, or stained for microscopy using either mouse anti-human CD40, mouse anti-human CD81 or mouse anti-human ICAM-1, followed by secondary staining using an anti-mouse AF647-conjugated monovalent nanobody (Chromotek). Cells were spotted and mounted on glass slides using SlowFade gold (Invitrogen). Slides were imaged in 3D on a Zeiss LSM 980 using the Airyscanning module and a 63X (1.4) oil objective. Using ImageJ, images were adjusted for brightness and contrast and max intensity projections were produced.

### PHA-stimulation of PBMCs

PBMCs were either fixed resting using 1% PFA, or were activated using 1.25 μg/ml PHA-L (Invitrogen) for 72h. Activated cells were harvested and fixed using 1% PFA. A fractions of the resting and activated cells were profiled using PNA, and a fraction were stained using anti-CD3 APC, anti-CD40 Pe, anti-CD19 AF700 and anti-HLA-DR Fitc antibodies and processed on a CytoFLEX (Beckman Coulter) and analyzed using FlowJo. Cells were gated for CD3+, CD19-for both the PNA and flow analysis.

Additionally, resting and activated cells were stained for microscopy using mouse anti-human CD40 (mIgG1) and mouse anti-human CD4 (mIgG2b) and an AF488-conjugated anti-mIgG2b nanobody and an AF647-conjugated anti-mIgG1 nanobody (Chromotek). Cells were spotted, mounted, imaged and analyzed according to the Raji microscopy protocol.

### Proximity Network Assay

Manufacturer’s instructions were followed for the Pixelgen Proxiome Kit. In short, samples were first fixed in 1% PFA, blocked, then stained overnight (16-20h) at +4C using a pool of 155 barcoded target specific antibodies and 3 mouse isotype controls, containing either type 1 or type 2 oligonucleotide sequence. Cells were then washed and antibody barcodes subjected to localized rolling circle amplification of gap-filled padlock probes for 10 minutes followed by addition of linker oligonucleotides designed to hybridize to two proximal RCA products of type 1 and 2. Gap-fill ligation was then performed to incorporate the UMI and PID sequences of the RCA products onto the hybridized linker oligonucleotide. One thousand cells were then transferred to a PCR amplification reaction to amplify the generated amplicons, followed by a second PCR reaction to attach sequencing adapters.

The PCR products were purified using SPRIselect beads, and quantified using Qubit hsDNA assay according to instructions of the Pixelgen Proxiome kit. The purified and quantified products were diluted to either 0.65 nM for Illumina P2 Xleap kits, or 0.49 nM for Illumina P3 and P4 Xleap kits and spiked with 15% PhiX and paired-end sequenced on an Illumina NextSeq2000 sequencing system, using 44 cycles for Read 1 and 78 cycles for Read 2.

### Data processing

PNA sequencing data were processed using Pixelator; an in-house open-source data analysis pipeline implemented in nextflow nf-core/pixelator (https://github.com/nf-core/pixelator)^10^. In short, sequenced reads undergo quality control and filtering before being assembled into amplicons representing the link between two proteins. Next, amplicons are grouped based on their marker assignments and then error-corrected to generate an edge list, which forms a graph of the entire sample. Within this comprehensive graph, individual cells are identified using Leiden community detection. Only those cells (graphs) exceeding a minimum size threshold of 8,000 nodes are retained. Finally, spatial information is calculated for each individual cellular component and stored, along with additional data, in a single PXL file. Downstream analyses were performed in R using pixelatorR v0.13.0 (https://github.com/PixelgenTechnologies/pixelatorR), pixelatorES v0.6.0 (https://github.com/PixelgenTechnologies/pixelatores), and Seurat v5.2.1. For graphical visualizations, ggplot2 v3.5.1 and ComplexHeatmap v2.22.0 were used.

### Cell Calling

Graph components corresponding to single cells were identified by inspecting Molecule Rank Plots (**Supplementary Figure 13A**), which are analogous to the Barcode Rank Plot used in scRNA-seq. A threshold is set at the size (number of nodes) corresponding to the point where rapid decline in size is observed, indicating that the distribution transitions from genuine single cell derived molecular graphs to partial components or debris. The input number of cells for library preparation is strongly correlated to the number of graph networks generated, indicating that graphs are generated from complete cells (**Supplementary Figure 13B**)

### Protein abundance

For each protein, abundance is quantified as the number of nodes in the component graphs whose antibody barcode (Ab-BC) has a Protein ID (PID) matching that protein. This count is computed separately for each component graph, where each component represents one cell, yielding a protein abundance value for every protein in every cell.

### UMAP visualization

The UMAP plots are based on the protein abundance of proteins that are expressed and variable in the visualized dataset. For the detailed cell type annotation of a PBMC sample in Figure 3, the dimensionality reduction was based on the intersection of 75 variable features identified by the Seurat function FindVariableFeatures and 58 manually curated lineage markers.

### Graph 3D layout visualization

Component graphs are projected in 2D or 3D space using the Weighted Pivot Multidimensional Scaling (wpmds) algorithm. For visualization purposes, individual nodes were colored according to the local density of the visualized marker(s) in a local neighbourhood around each node using the Getis-Ord local statistics or by smoothing abundance levels over their neighbours in Euclidean space.

### Join Count statistics

The Proximity Score was used to assess the degree of association between two markers in a component graph, and is based on join count statistics^22^. The developed algorithm tracks the number of edges between marker pairs in a component, called “join count”, corresponding to the number of times instances of the two marker species were adjacent to each other. In order to understand whether the join count was higher or lower compared to chance, a baseline expectation was calculated using Monte Carlo simulations, where the spatial distribution of markers was randomized while the graph structure and marker abundance were kept constant. For each graph component (corresponding to a single cell) 100 such permutations were created, forming a null distribution that describes the join count of pairs of markers if they were randomly placed as node attributes on the graph, enabling comparison of the observed patterns to random expectation.

Using the mean (*E*) and standard deviation (*σ*) of the simulated null distribution, the join counts (*x*) were formulated as a log2 ratio (L):

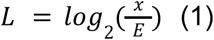

And a Z-score (Z):

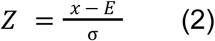

Formulating the join count as a log2 ratio or a Z-score relating it to random expectation effectively represents an abundance normalization of the score. Highly abundant protein pairs can show lower Proximity Scores, as the abundance-driven random chance colocalization is higher for abundant protein pairs.

### Marker selection for Proximity Score visualizations

For visualizations of Proximity Scores in e.g. colocalization heatmaps and networks, proteins were selected by passing a minimum abundance and frequency within the specific cell population. Isotype controls and background levels within negative populations were used to set the cut-offs.

### Patch detection

Patches were detected by expanding the adjacency matrix of a component to include the second neighbors followed by subsetting this expanded adjacency matrix to nodes that are labeled by B-cell originating patch-specific markers CD20, CD22 and CD40 as well as the nodes connecting these patch-specific marker nodes. A new graph was created from the subsetted adjacency matrix and the graph was split into its connected components representing the different patches. Next, Leiden iterative community detection was used to split up weakly connected patch components (resolution parameter = 0.01) and patches smaller than 200 nodes were filtered out. To ensure that the results were not biased by the choice of threshold, the analysis was repeated twice, filtering patches smaller than 100 and 400 nodes, respectively. (**Supplementary Figure 9D**).

---

**Supplementary Figure 1.**
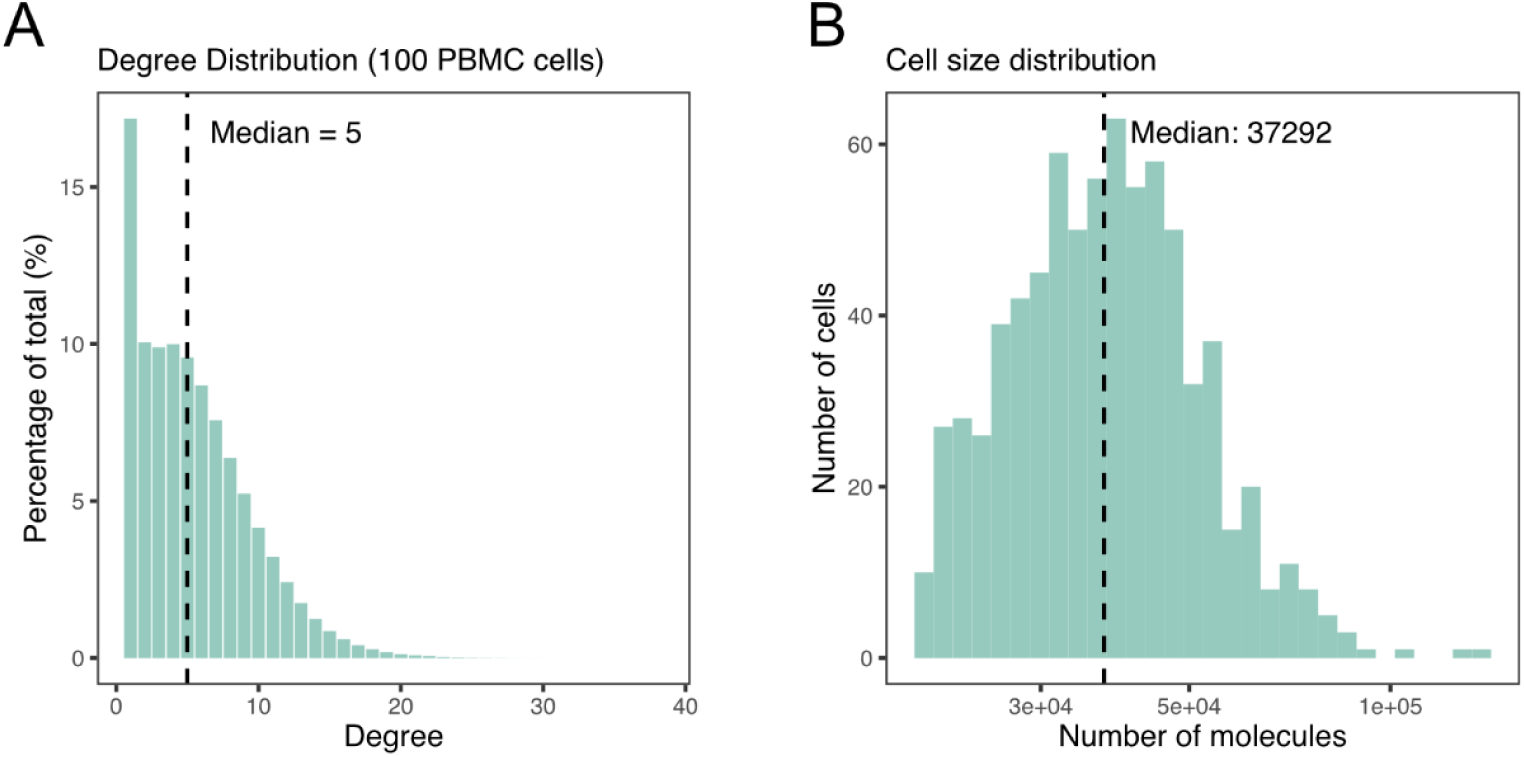
Number of spatial nodes and connections per PNA protein. (A) Histogram showing the distribution of node degree across PNA nodes in 100 random cells for PBMC cells. The dashed line indicates the median. (B) Histogram of cell sizes across the PBMC sample. Size is here defined as the number of uniquely positioned protein molecules (nodes) in the cell graph. The dashed line indicates the median.

**Supplementary Figure 2.**
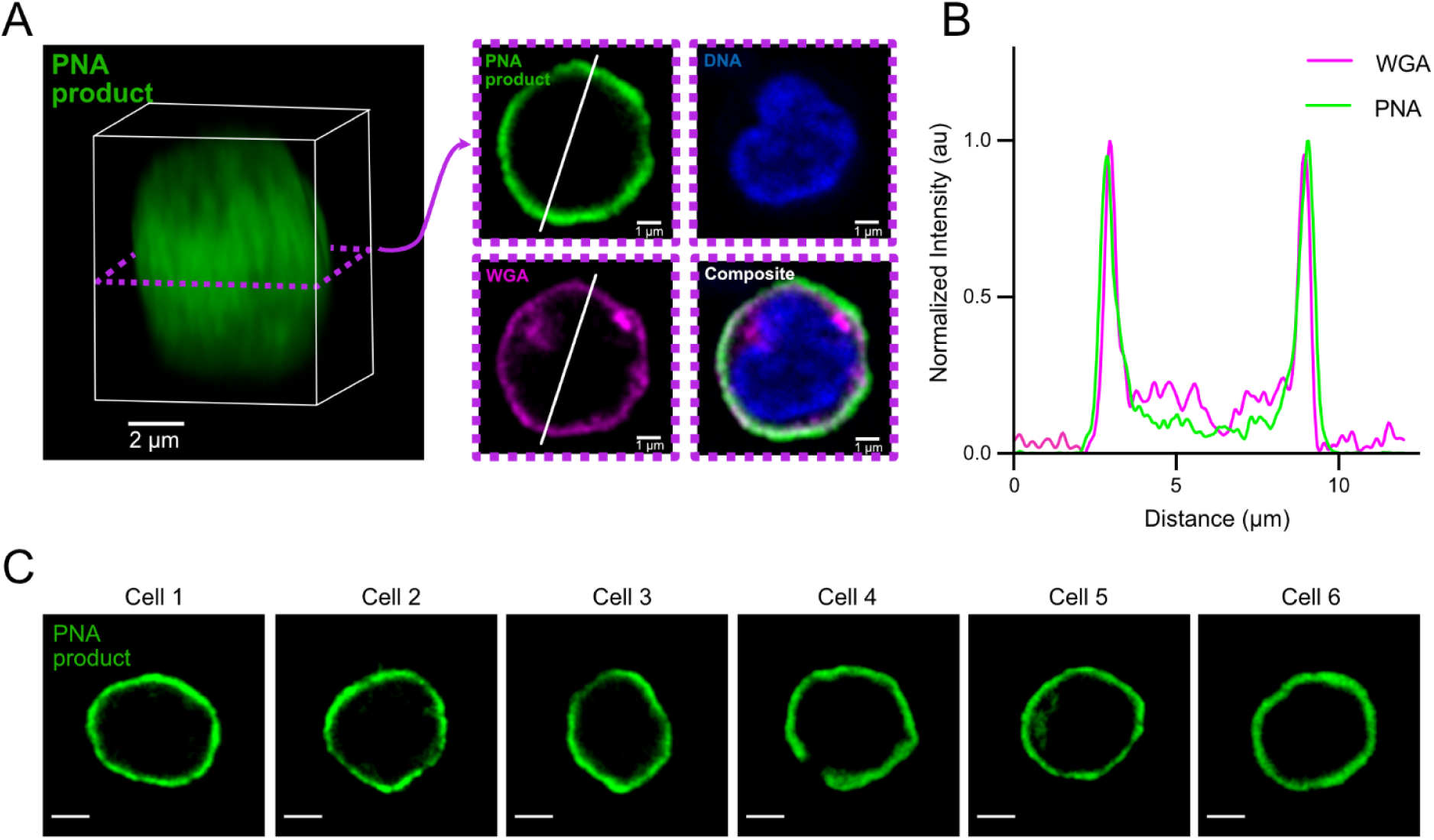
Distribution of PNA reaction products at the cell surface. (A) Fluorescence microscopy image of PNA products surrounding a single PBMC immune cell. The RCA reaction was supplemented with fluorescently labelled nucleotides to visualize the cell surface coverage with PNA products. Left, 3D reconstruction; Right, single imaging plane. (B) Fluorescence intensity profiles for WGA (labelling the plasma membrane) and PNA products along the line indicated in (A), demonstrating specific localization of the product to the cell membrane. (C) Single imaging plane of the fluorescent PNA product in 6 randomly selected PBMCs. Scale bars: 2 μm.

**Supplementary Figure 3:**
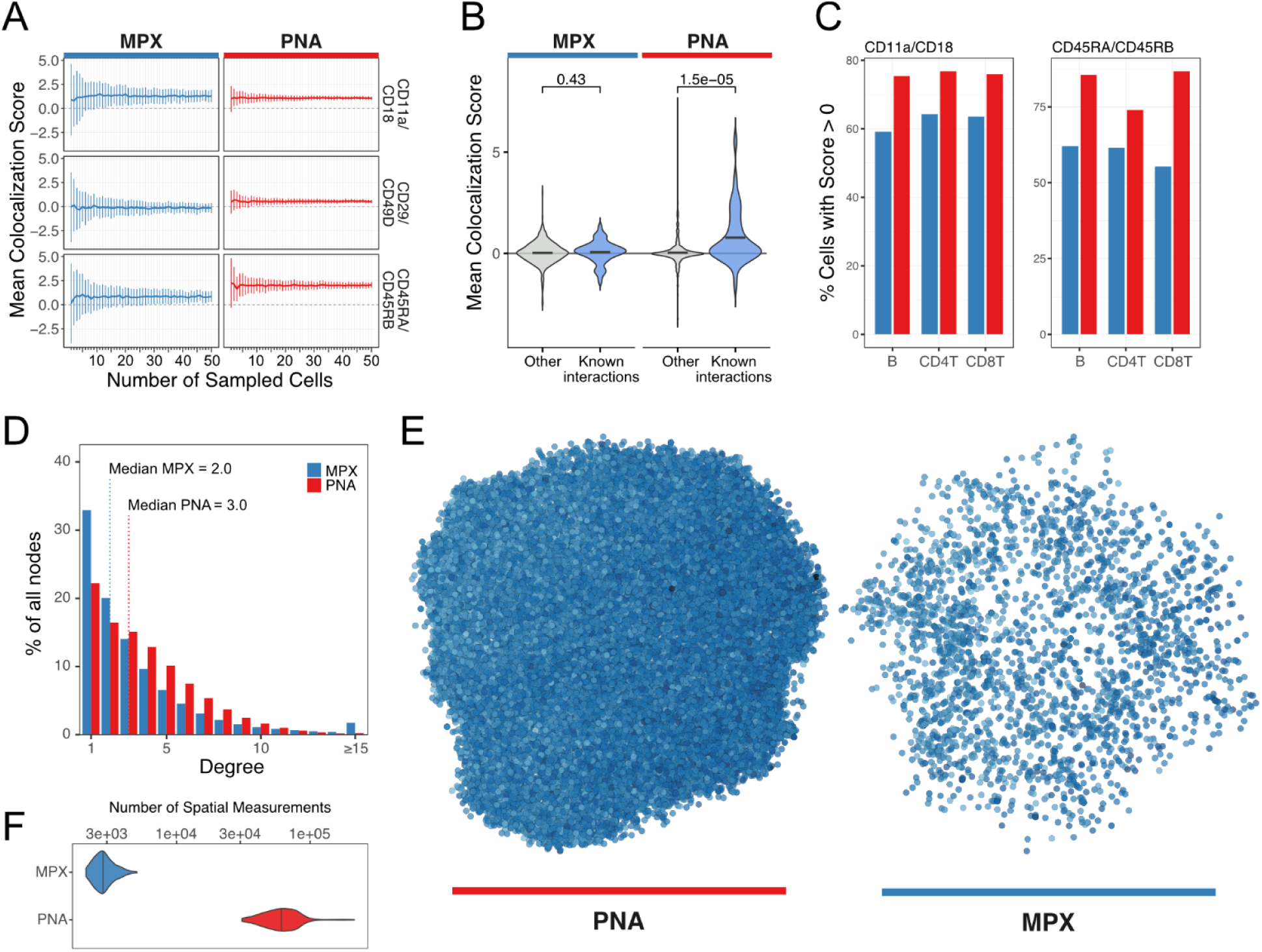
Comparison of PNA technology to MPX. (A) Comparison of mean colocalization scores for three established interaction pairs in CD8 T cells, contrasting MPX and PNA technologies. The data presented are mean colocalization scores with standard deviation, derived from 100 sampling iterations. (B) Assessment of mean colocalization scores between known interactions (sourced from BioGrid Release 5.0.254 [Oughtred *et al.*, *Protein Sci.* (2021)]) and the other pairs in CD8 T cells, comparing MPX and PNA. The significance level for the Wilcoxon test between these two sets is shown. (C) Percentage of CD8 T cells exhibiting a positive colocalization score for known interacting pairs across various cell populations. (D) Degree distribution of nodes across 100 CD8 T cells in MPX and PNA. (E) Representative spatial layouts of single CD8 T cells as determined by MPX and PNA. (F) Summary of the number of spatial measurements per technology in a CD8 T cell population, showing the number of pixels for MPX and the number of molecules for PNA.

**Supplementary Figure 4.**
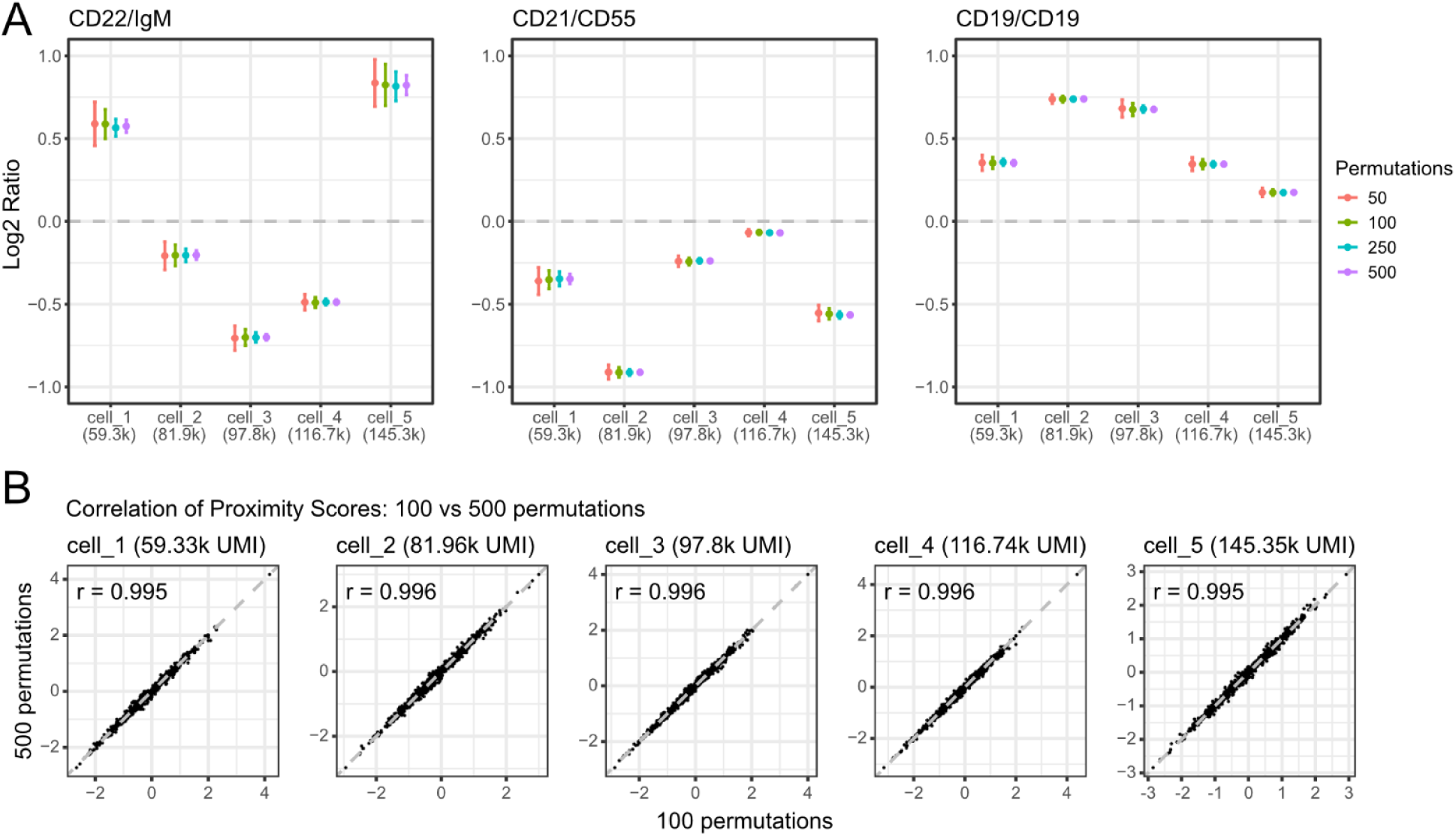
Characterization of the Proximity Score. (A) log_2_ ratio Proximity Scores of selected protein pairs (CD22/IgM, CD21/CD55, and CD19/CD19) for 5 cells across a range of sizes and number of permutations. The point indicates the mean log_2_ ratio Proximity Score of 100 repeated Proximity Score calculations and the whiskers indicate 1 standard deviation. (B) Scatterplot showing the correlation of log_2_ ratio Proximity Scores when using 100 permutations (x-axis) versus 500 permutations (y-axis) to construct the null distribution per each cell.

**Supplementary Figure 5.**
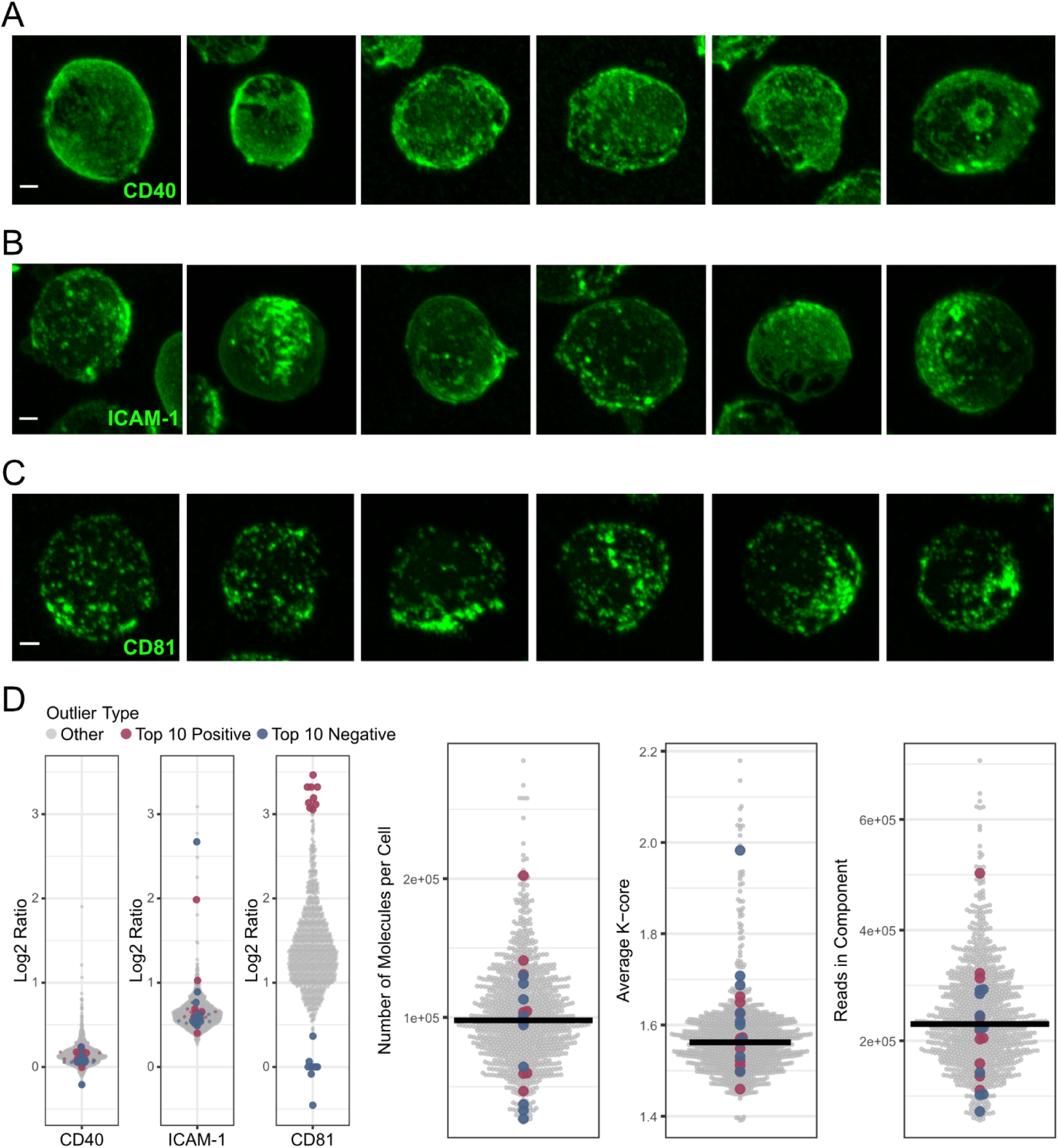
(A-C) Fluorescent microscopy images of 18 randomly selected Raji cells stained for CD40 (A), ICAM-1 (B) and CD81 (C). Scale bars: 2 μm. (D) Characterization of the Raji cells with top 10 highest and top 10 lowest CD81 Proximity Scores. Shown are the log_2_ ratio Proximity Scores of CD40, CD54 (ICAM-1), and CD81, as well as QC metrics; number of proteins, average k-core across nodes, and number of reads in the component.

**Supplementary Figure 6.**
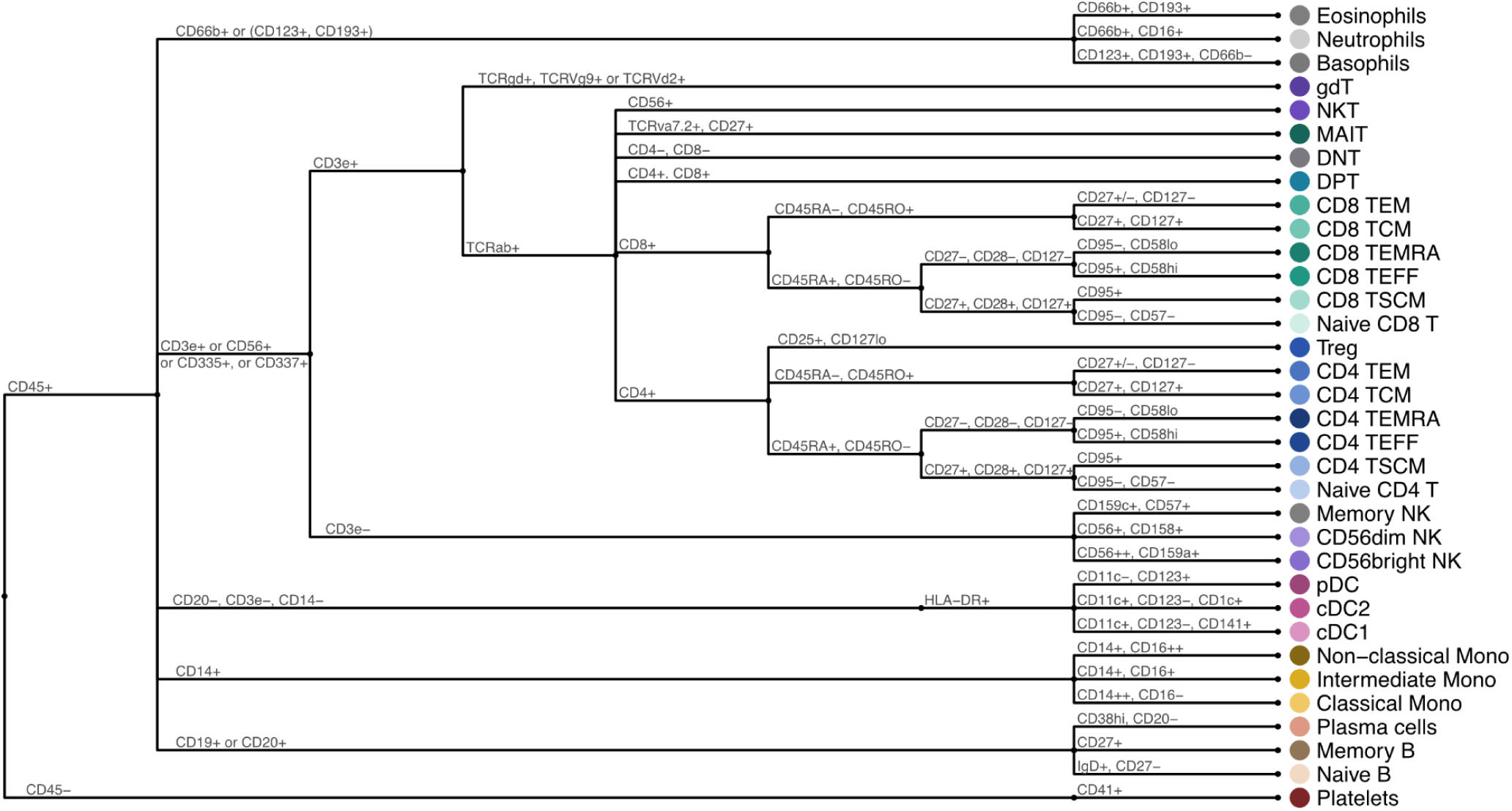
Hierarchical annotation strategy of cell types. TSCM, stem memory T cells; TCM, central memory T cells; TEM, effector memory T cells; TEFF, effector T cells; TEMRA, effector memory re-expressing CD45RA T cells; Treg, regulatory T cells; MAIT, mucosal-associated invariant T cells; DNT double-negative T lymphocytes.

**Supplementary Figure 7.**
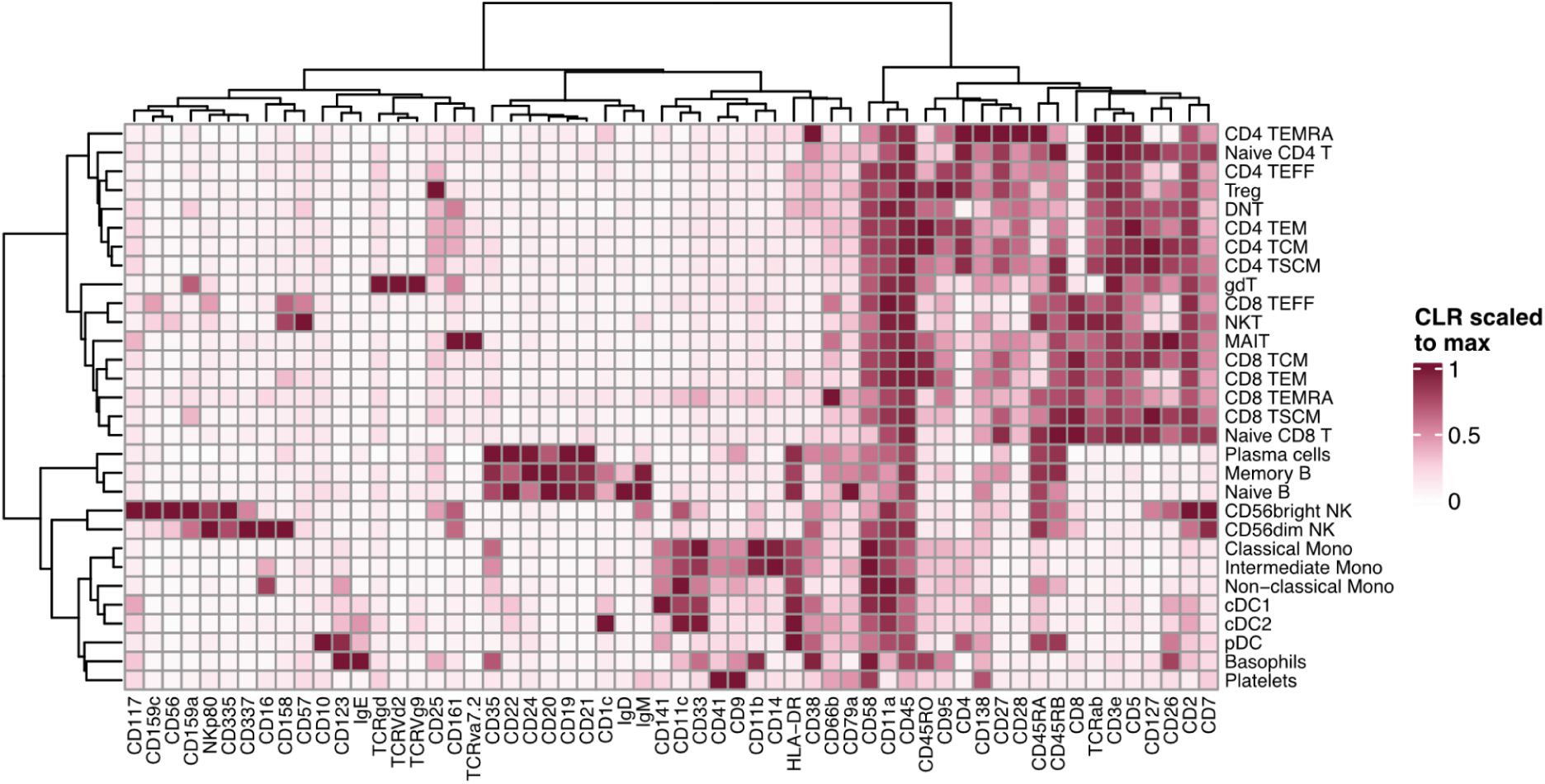
Heatmap of relative mean abundance level across annotated cell types in PBMCs. The mean abundance (CLR) has here been max-scaled such that the population with the highest abundance has a value of 1. TSCM, stem memory T cells; TCM, central memory T cells; TEM, effector memory T cells; TEFF, effector T cells; TEMRA, effector memory re-expressing CD45RA T cells; Treg, regulatory T cells; MAIT, mucosal-associated invariant T cells; DNT double-negative T lymphocytes.

**Supplementary Figure 8.**
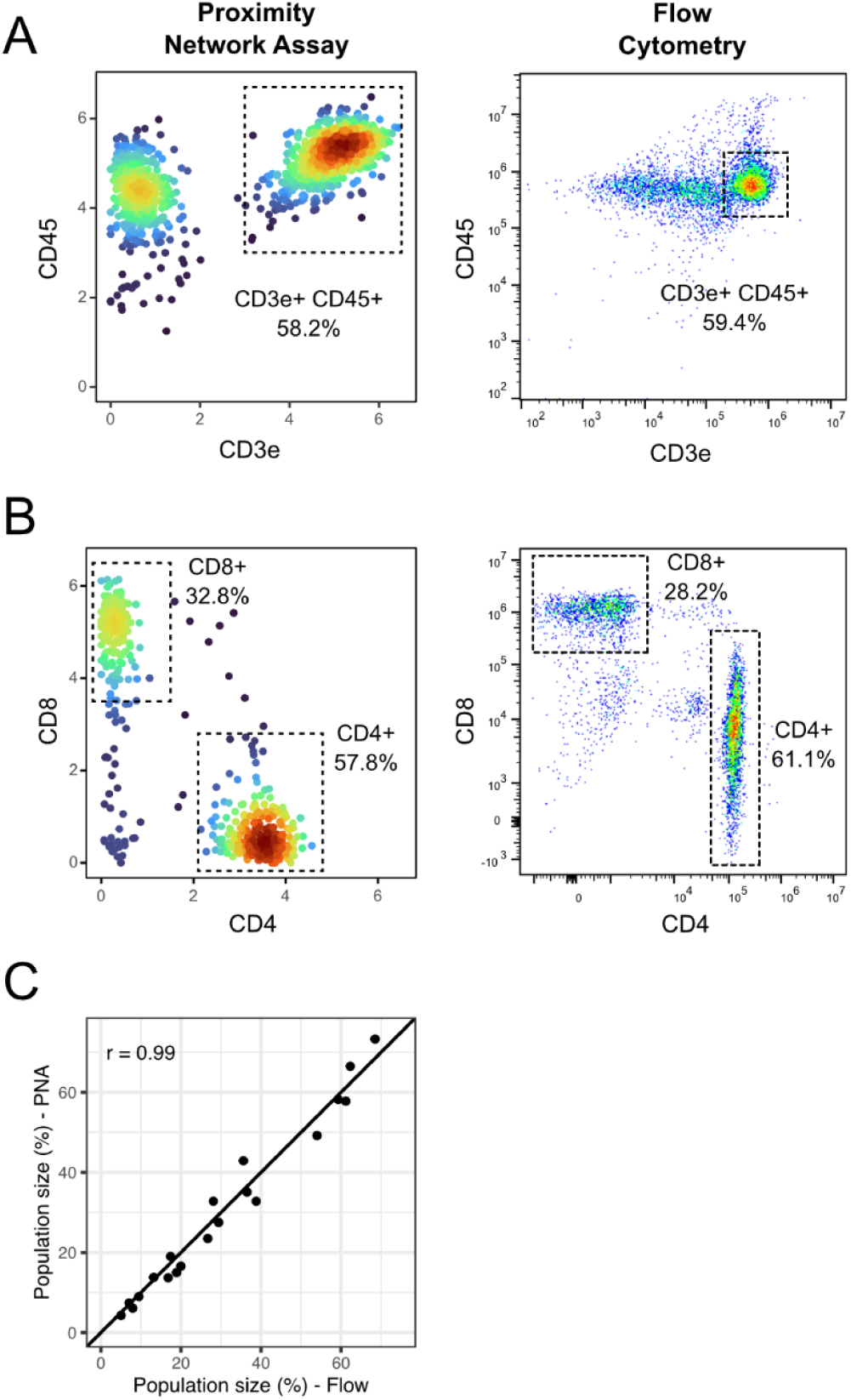
Comparison of the Proximity Network Assay and Flow Cytometry for cell identification. (A) Gating of T cells using density plots for CD3e and CD45 for PNA (left) and flow cytometry (right). (B) Gating of CD4 T cells (CD4^+^, CD8^-^) and CD8 T cells (CD8^+^, CD4^-^) for PNA (left) and flow cytometry (right). (C) Scatter plot showing the correlation between PNA and flow cytometry for the detection of up to 8 unique cell populations in 3 different biological donors (20 identified populations).

**Supplementary Figure 9.**
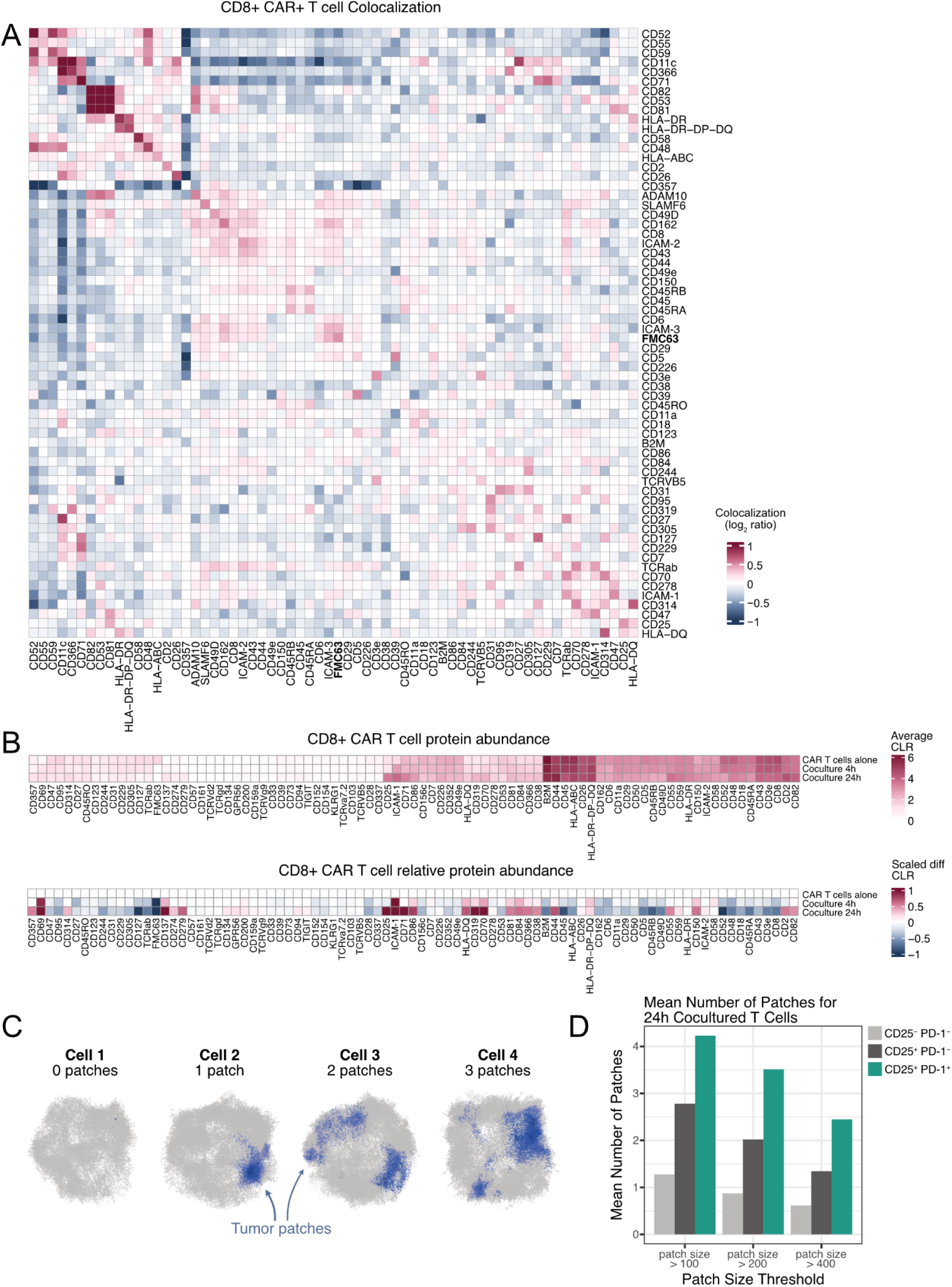
Proximity Network of CD19 CAR-T cells. (A) Heatmap of mean Proximity Scores (log2 ratio) of abundant proteins in CD8^+^ FMC63+ CAR-T cells showing protein colocalization and self-clustering. (B) Average protein abundance of CD8+ T cells alone, at 4h and at 24h of coculture (top panel) and scaled differential abundance of the cocultured T cells at 4h or 24h as compared to T cells alone (bottom panel). The Axis has been capped at +6 (top panel) and −1 and 1 (bottom panel). (C) Representative 3D visualizations of 4 different PNA cell graphs containing various amounts of Raji-derived membrane patches (the patches consist of CD40, CD20, CD22 and IgM counts). (D) Effect of patch size threshold on mean number of patches on 24h cocultured CD8+ T cells.

**Supplementary Figure 10.**
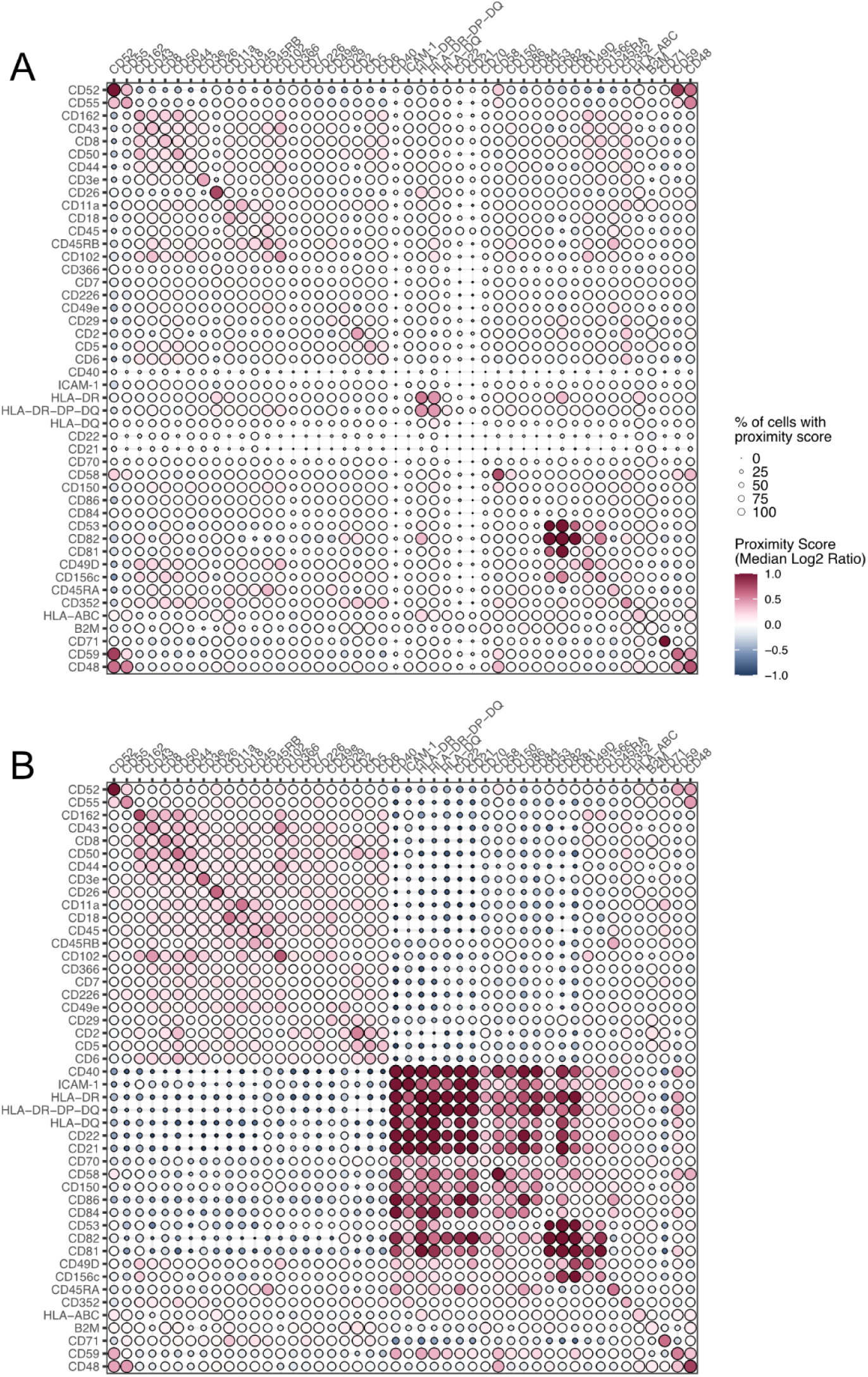
Colocalization heatmaps of unstimulated (A) and 24h cocultured (B) CAR T cells, displaying the acquisition of a protein domain following coculture.

**Supplementary Figure 11.**
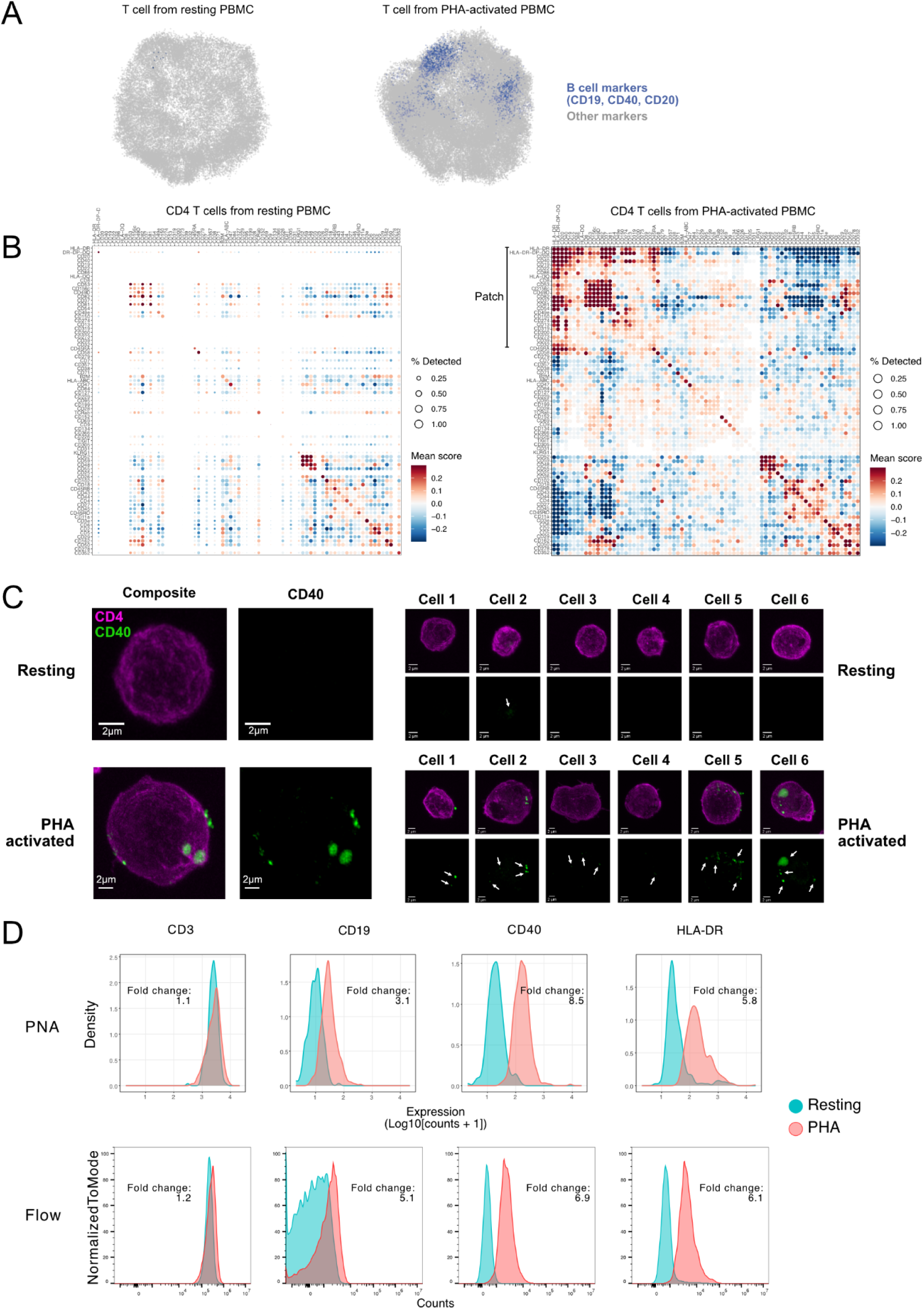
Trogocytosis in T cells from PHA-activated PBMCs. (A) Representative 3D visualizations of PNA graphs of a resting T cell (left) and a PHA-activated T cell with B cell patches (right). (B) Heatmap of mean Proximity Scores (log2 ratio) of resting CD4+ T cells (left) and PHA-activated CD4+ T cells (right) showing the acquisition of a B cell-derived membrane domain. (C) Microscopy validation of CD40 trogocytosis in 7 randomly selected resting and 7 randomly selected PHA-activated CD4 T cells. Scale bars: 2 μm. (D) Validation of trogocytosis through the assessment of protein abundance of CD3 (control), CD19, CD40 and HLA-DR in resting and PHA-activated CD3+ T cells as measured by PNA (top) and flow cytometry (bottom).

**Supplementary Figure 12.**
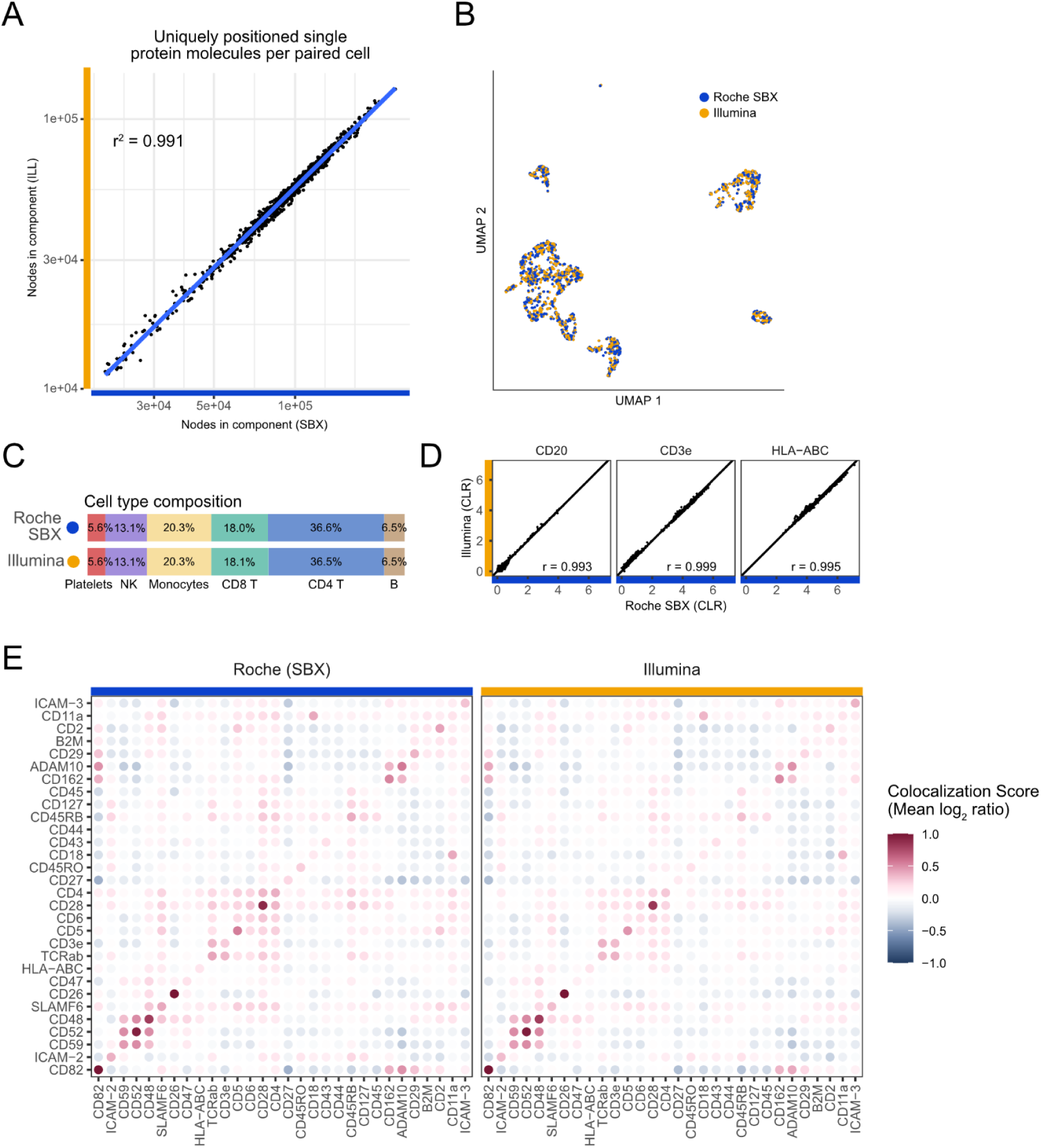
High throughput nanopore sequencing compared to sequencing by synthesis. Two aliquots of the same Proximity Network Assay library were subjected to sequencing by Illumina SBS and Roche SBX. (A) Scatterplot of uniquely identified proteins per paired single cells compared across platforms. (B) UMAP of cells identified in data generated by either platform. (C) Barplot comparing annotated cell type composition between the platforms. (D) Scatterplots comparing abundance (CLR) of three selected markers (CD20, CD3e, and HLA-ABC) between paired cells generated by Illumina SBS and Roche SBX. E) Protein colocalization heatmaps showing mean log2 ratio Proximity Scores of CD4 T cells for Roche SBX (left) and Illumina SBS (right).

**Supplementary Figure 13.**
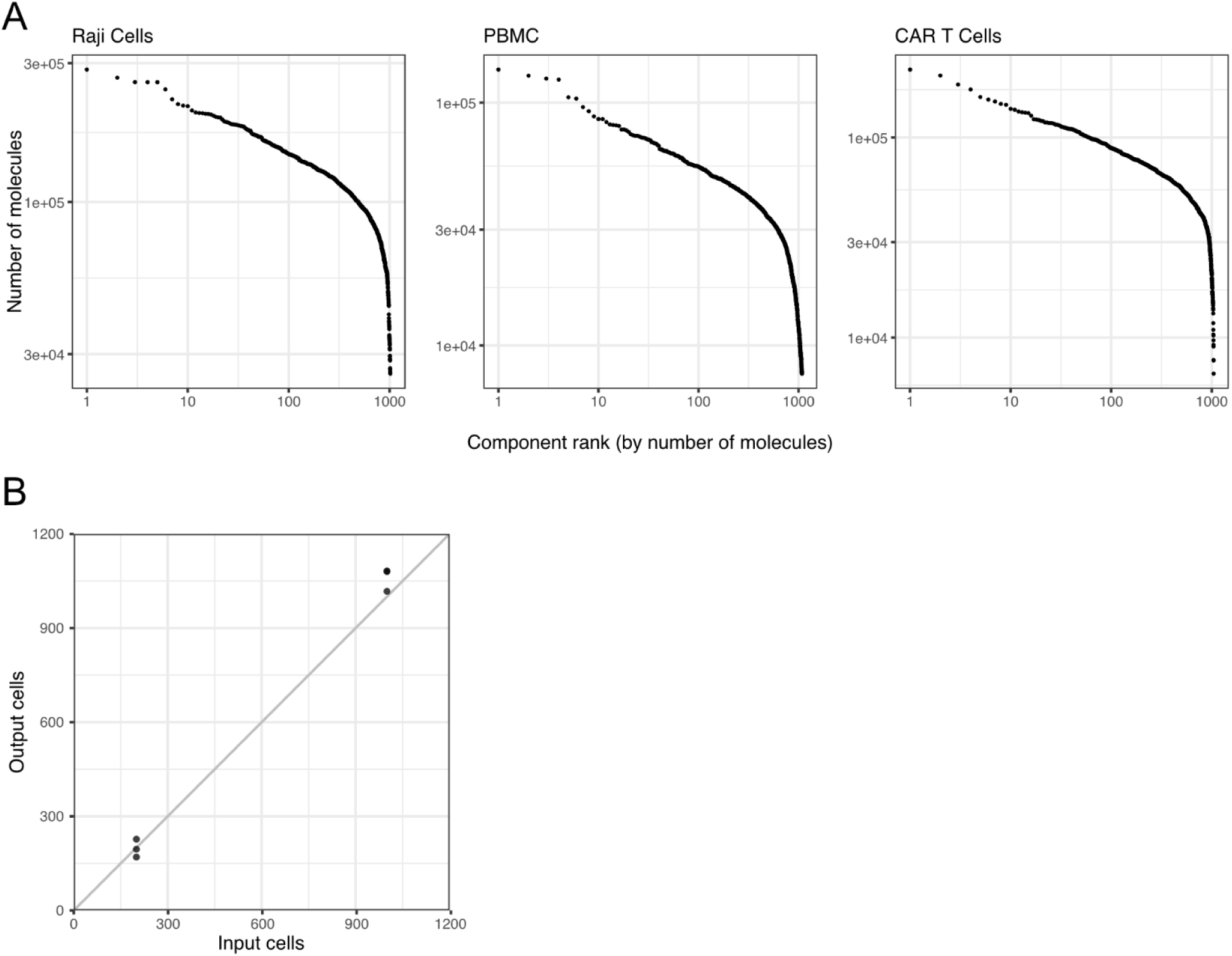
Cell calling in PNA. (A) Molecule Rank Plots showing the distribution of graph component sizes for samples of Raji cells (left), PBMC (middle), and CAR T cells (right). The rapidly declining size is an indicator that the distribution transitions from real cell graphs to debris or disconnected partial graphs, and can be used as a basis of cell filtering. (B) Scatterplot showing the input cells versus the called output cells in the PNA data for 3 samples of 200 cells and 3 samples of 1000 cells.

## Notes

### Summary of Updates

Title changed to “Cells as Single-Molecule Protein Proximity Networks.” Interactomics language toned down throughout; the method is now framed as measuring protein proximity and membrane organization. Added orthogonal validation by flow cytometry and microscopy (Supplementary Figures 2, 5, 8, 11) and a side-by-side benchmark of PNA versus Molecular Pixelation (Supplementary Figure 3). Added Supplementary Figure 1 (link-count and cell-size distributions), Supplementary Figure 4 (Proximity Score robustness across permutations and cell sizes), Supplementary Figure 9 (trogocytosis patch cutoff robustness), and Supplementary Figure 13 (molecule rank plot and cell-input versus graph-output correlation). Figure 1 and Methods updated to clarify the method, cell calling, type-1/type-2 antibody design, and protein abundance inference. Figure 5 (SLE cohort) expanded; CAR-T trogocytosis analysis strengthened. Added nanopore sequencing of PNA libraries (Roche SBX) for a fully optics-free workflow, plus a supplementary video of the 3D Proximity Network data.

## References

1. Al-Aghbar MA, Jainarayanan AK, Dustin ML, Roffler SR. The interplay between membrane topology and mechanical forces in regulating T cell receptor activity. Commun Biol. 2022;5(1):40. doi:10.1038/s42003-021-02995-1

2. Dustin ML. The Immunological Synapse. Cancer Immunol Res. 2014;2(11):1023–1033. doi:10.1158/2326-6066.CIR-14-0161

3. Viganò S, Utzschneider DT, Perreau M, Pantaleo G, Zehn D, Harari A. Functional Avidity: A Measure to Predict the Efficacy of Effector T Cells? Clin Dev Immunol. 2012;2012:1–14. doi:10.1155/2012/153863

4. Manes S, Gomezmouton C, Lacalle R, Jimenezbaranda S, Mira E, Martineza C. Mastering time and space: immune cell polarization and chemotaxis. Semin Immunol. 2005;17(1):77–86. doi:10.1016/j.smim.2004.09.005

5. Lu H, Zhou Q, He J, et al. Recent advances in the development of protein–protein interactions modulators: mechanisms and clinical trials. Signal Transduct Target Ther. 2020;5(1):213. doi:10.1038/s41392-020-00315-3

6. Rudnicka D, Oszmiana A, Finch DK, et al. Rituximab causes a polarization of B cells that augments its therapeutic function in NK-cell-mediated antibody-dependent cellular cytotoxicity. Blood. 2013;121(23):4694–4702. doi:10.1182/blood-2013-02-482570

7. Söderberg O, Gullberg M, Jarvius M, et al. Direct observation of individual endogenous protein complexes in situ by proximity ligation. Nat Methods. 2006;3(12):995–1000. doi:10.1038/nmeth947

8. Vistain L, Van Phan H, Keisham B, et al. Quantification of extracellular proteins, protein complexes and mRNAs in single cells by proximity sequencing. Nat Methods. 2022;19(12):1578–1589. doi:10.1038/s41592-022-01684-z

9. Chandrasekharan G, Unnikrishnan M. High throughput methods to study protein-protein interactions during host-pathogen interactions. Eur J Cell Biol. 2024;103(2):151393. doi:10.1016/j.ejcb.2024.151393

10. Karlsson F, Kallas T, Thiagarajan D, et al. Molecular pixelation: spatial proteomics of single cells by sequencing. Nat Methods. 2024;21(6):1044–1052. doi:10.1038/s41592-024-02268-9

11. Boulgakov AA, Ellington AD, Marcotte EM. Bringing Microscopy-By-Sequencing into View. Trends Biotechnol. 2020;38(2):154–162. doi:10.1016/j.tibtech.2019.06.001

12. Hoffecker IT, Yang Y, Bernardinelli G, Orponen P, Högberg B. A computational framework for DNA sequencing microscopy. Proc Natl Acad Sci U S A. 2019;116(39):19282–19287. doi:10.1073/pnas.1821178116

13. Weinstein JA, Regev A, Zhang F. DNA Microscopy: Optics-free Spatio-genetic Imaging by a Stand-Alone Chemical Reaction. Cell. 2019;178(1):229–241.e16. doi:10.1016/j.cell.2019.05.019

14. Abdulraouf A, Jiang W, Xu Z, et al. Optics-free Spatial Genomics for Mapping Mouse Brain Aging. Genomics. Preprint posted online August 8, 2024. doi:10.1101/2024.08.06.606712

15. Liao H, Kottapalli S, Huang Y, et al. Optics-free reconstruction of 2D images via DNA barcode proximity graphs. BioRxiv Prepr Serv Biol. Published online August 8, 2024:2024.08.06.606834. doi:10.1101/2024.08.06.606834

16. Qian N, Weinstein JA. Spatial transcriptomic imaging of an intact organism using volumetric DNA microscopy. Nat Biotechnol. 2026;44(2):222–232. doi:10.1038/s41587-025-02613-z

17. Fredriksson S, Dixon W, Ji H, Koong AC, Mindrinos M, Davis RW. Multiplexed protein detection by proximity ligation for cancer biomarker validation. Nat Methods. 2007;4(4):327–329. doi:10.1038/nmeth1020

18. Lundberg M, Eriksson A, Tran B, Assarsson E, Fredriksson S. Homogeneous antibody-based proximity extension assays provide sensitive and specific detection of low-abundant proteins in human blood. Nucleic Acids Res. 2011;39(15):e102–e102. doi:10.1093/nar/gkr424

19. Feng W, Beer JC, Hao Q, et al. NULISA: a proteomic liquid biopsy platform with attomolar sensitivity and high multiplexing. Nat Commun. 2023;14(1):7238. doi:10.1038/s41467-023-42834-x

20. Majstoravich S, Zhang J, Nicholson-Dykstra S, et al. Lymphocyte microvilli are dynamic, actin-dependent structures that do not require Wiskott-Aldrich syndrome protein (WASp) for their morphology. Blood. 2004;104(5):1396–1403. doi:10.1182/blood-2004-02-0437

21. Zhong L, Chen H, Cao S, Hu S. Single Nucleotide Recognition and Mutation Site Sequencing Based on a Barcode Assay and Rolling Circle Amplification. Biosensors. 2024;14(11):521. doi:10.3390/bios14110521

22. Moran PAP. The Interpretation of Statistical Maps. J R Stat Soc Ser B Stat Methodol. 1948;10(2):243–251. doi:10.1111/j.2517-6161.1948.tb00012.x

23. Pavlasova G, Mraz M. The regulation and function of CD20: an “enigma” of B-cell biology and targeted therapy. Haematologica. 2020;105(6):1494–1506. doi:10.3324/haematol.2019.243543

24. Loertscher R, Lavery P. The role of glycosyl phosphatidyl inositol (GPI)-anchored cell surface proteins in T-cell activation. Transpl Immunol. 2002;9(2):93–96. doi:10.1016/S0966-3274(02)00013-8

25. Arp AB, Abel Gutierrez A, Ter Beest M, et al. CD70 recruitment to the immunological synapse is dependent on CD20 in B cells. Proc Natl Acad Sci. 2025;122(16):e2414002122. doi:10.1073/pnas.2414002122

26. Wentink MWJ, van Zelm MC, van Dongen JJM, Warnatz K, van der Burg M. Deficiencies in the CD19 complex. Clin Immunol. 2018;195:82–87. doi:10.1016/j.clim.2018.07.017

27. Cherukuri A, Cheng PC, Pierce SK. The Role of the CD19/CD21 Complex in B Cell Processing and Presentation of Complement-Tagged Antigens. J Immunol. 2001;167(1):163–172. doi:10.4049/jimmunol.167.1.163

28. Garcia-Parajo MF, Cambi A, Torreno-Pina JA, Thompson N, Jacobson K. Nanoclustering as a dominant feature of plasma membrane organization. J Cell Sci. 2014;127(23):4995–5005. doi:10.1242/jcs.146340

29. Schmidt TH, Homsi Y, Lang T. Oligomerization of the Tetraspanin CD81 via the Flexibility of Its δ-Loop. Biophys J. 2016;110(11):2463–2474. doi:10.1016/j.bpj.2016.05.003

30. Hamieh M, Dobrin A, Cabriolu A, et al. CAR T cell trogocytosis and cooperative killing regulate tumour antigen escape. Nature. 2019;568(7750):112–116. doi:10.1038/s41586-019-1054-1

31. Olson ML, Mause ERV, Radhakrishnan SV, et al. Low-affinity CAR T cells exhibit reduced trogocytosis, preventing rapid antigen loss, and increasing CAR T cell expansion. Leukemia. 2022;36(7):1943–1946. doi:10.1038/s41375-022-01585-2

32. Gjertsson I, McGrath S, Grimstad K, et al. A close-up on the expanding landscape of CD21-/low B cells in humans. Clin Exp Immunol. 2022;210(3):217–229. doi:10.1093/cei/uxac103

33. Koarada S, Tada Y, Ushiyama O, et al. B cells lacking RP105, a novel B cell antigen, in systemic lupus erythematosus. Arthritis Rheum. 1999;42(12):2593–2600. doi:10.1002/1529-0131(199912)42:12%3C2593::AID-ANR12%3E3.0.CO;2-G

34. You M, Dong G, Li F, et al. Ligation of CD180 inhibits IFN-α signaling in a Lyn-PI3K-BTK-dependent manner in B cells. Cell Mol Immunol. 2017;14(2):192–202. doi:10.1038/cmi.2015.61

35. Carlos AJ, Yang D, Thomas DM, Huang S, Harter KI, Moellering RE. Family-Wide Photoproximity Profiling of Integrin Protein Social Networks in Cancer. Cancer Biology. Preprint posted online September 22, 2024. doi:10.1101/2024.09.18.613588

36. Floyd BM, Schmidt EL, Till NA, et al. Mapping the nanoscale organization of the human cell surface proteome reveals new functional associations and surface antigen clusters. Systems Biology. Preprint posted online February 17, 2025. doi:10.1101/2025.02.12.637979

37. Naei VY, Tubelleza R, Monkman J, et al. Spatial interaction mapping of PD-1/PD-L1 in Head and Neck Cancer reveals the role of Macrophage-Tumour Barriers associated with immunotherapy response. In Review. Preprint posted online November 7, 2024. doi:10.21203/rs.3.rs-5398442/v2

38. Cesnik A, Takacsi-Nagy O, Le T, Roth TL, Satpathy AT, Lundberg E. Molecular pixelation of the CAR T cell surface proteome. Systems Biology. Preprint posted online February 2, 2026. doi:10.64898/2026.01.30.702970

39. Sezgin E, Levental I, Mayor S, Eggeling C. The mystery of membrane organization: composition, regulation and physiological relevance of lipid rafts. Nat Rev Mol Cell Biol. 2017;18(6):361–374. doi:10.1038/nrm.2017.16

40. Levy S, Shoham T. The tetraspanin web modulates immune-signalling complexes. Nat Rev Immunol. 2005;5(2):136–148. doi:10.1038/nri1548

41. Till NA, Ramanathan M, Bertozzi CR. Induced proximity at the cell surface. Nat Biotechnol. 2025;43(5):702–711. doi:10.1038/s41587-025-02592-1

42. Cai S, Hu T, Venkataraman A, et al. Spatially resolved subcellular protein–protein interactomics in drug-perturbed lung-cancer cultures and tissues. Nat Biomed Eng. 2025;9(7):1039–1061. doi:10.1038/s41551-024-01271-x

43. Kokoris M, McRuer R, Nabavi M, et al. Sequencing by Expansion (SBX) – a novel, high-throughput single-molecule sequencing technology. Genomics. Preprint posted online February 24, 2025. doi:10.1101/2025.02.19.639056

